# Back to the Future: Environmental genomic selection to take advantage of polygenic local adaptation

**DOI:** 10.1101/2024.10.09.617488

**Authors:** Anna Halpin-McCormick, Quinn Campbell, Sónia Negrão, Peter L. Morrell, Sariel Hubner, Jeff Neyhart, Michael Kantar

## Abstract

The genetic basis of adaptation is a fundamental question in evolutionary genetics. Environmental association analysis (EAA) and various allele frequency comparisons in genomic environmental association (GEA) have become standard approaches for investigating the genetic basis of adaptation to natural environments. While these analyses provide insight into local adaptation, they have not been widely adopted in breeding or conservation programs. This may be attributable to the difficulty in identifying the best individuals for transplantation/relocation in conservation efforts or identification of the best parents in breeding programs. To explore the use of EAA and GEA for future breeding programs, we used a cereal crop - barley (*Hordeum vulgare* L.) as our case-study species due to its wide adaptability to different environments and agro-ecologies, ranging from marginal and low input fields to high-productive farms. Here, we use publicly available data to conduct environmental genomic selection (EGS) on 753 landrace barley accessions using a mini-core of 31 landrace accessions and a de-novo core of 100 as the training populations. Environmental genomic selection is to environmental association analysis (EAA) what genomic selection is to genome-wide association studies (GWAS). Since local adaptation to the environment is polygenic, a whole-genome approach is likely to be more accurate for selecting for environmental adaptation. Here we show distinct genetic background and population differences and how an integrative approach coupling environmental genomic selection and species distribution modelling can help identify key parents for breeding for adaptation to specific environmental variables and geographies to minimize linkage drag.

## Main

Climate resilience will require adaptation to changing climatic regimes, including warmer day and nighttime temperatures and more sporadic but potentially more intense rainfall events. For grassland and forest environments, this could cause major changes in species composition. In some cases, human efforts may be necessary to identify and relocate individuals or genotypes well adapted for these new environments. Agriculture faces a similar issue, in terms of a realignment of where crops are produced and which genetic backgrounds will be successful in new growth environments. Adapting to a changing environment is a major concern for natural and managed plant populations, this includes forests, orchards, and field crops (1,2). There is an expanding scope of research exploring methods for accelerating plant breeding for a changing climate (3,4). Many have emphasized an understanding of climate resilience targets to help define interventions that will be the most useful for specific geographies and species (5–8).

Early efforts to identify the genetic basis of environmental adaptation examined allele frequency differences among populations (9). Refinements to these approaches have sought to account for the variance among related individuals and populations relative to environmental variables (10,11) and to identify sharp changes in allele frequency in species with an essentially continuous geographic range (12). Collectively the approaches have been identified as landscape genomics (13–15), which uses evolutionary relationships among wild relatives of crops and landraces to identify potentially adaptive loci (7,16,17). A popular approach in landscape genomics involves genome-environment association (GEA) analysis, which seeks to identify relationships between allele frequencies or marker allele dosages and environmental variables that characterize the climate where each accession originates. Some early GEA studies made use of populations sampled from natural environments (18,19). Another popular source of samples for GEA has been germplasm collections where collection localities are represented by individual (often inbred) accessions and no population-level information is available. GEA studies often focus on identifying loci underlying local adaptation (20–22). The implicit assumption is that environmental variables such as temperature can be used as proxies for the selective pressures that have shaped the differences in allele frequencies among populations (23,24). By correlating genetic information with environmental data that is most relevant to an important abiotic stress in a crop (e.g., temperature for heat stress), GEA analyses provide a means to select individuals from germplasm collections that may confer tolerance to those stresses.

There are many frameworks for GEA analysis (25,26), however, in crops popular approaches involve the use of linear mixed-models, a well-established tool for other genomics-enabled plant breeding as well as quantitative genetic methods such as genome-wide association studies (GWAS) and genomic selection (GS). In many GEA analyses, the standard GWAS model is modified to use explicit bioclimatic data instead of trait phenotypic values as the response variable, in what is known as environmental GWAS (E-GWAS) (27). While standard GWAS has become an important method for understanding the genetic architecture of complex traits in crops, it is less useful as a plant breeding selection tool, since only a fraction of heritable polygenic variation may be detected (28,29). Instead, many breeding programs now use GS, which estimates the effect of genome wide markers simultaneously and predicts the breeding value or total genetic merit of individuals (30,31). Similarly, while E-GWAS can advance our understanding of the genetic architecture underlying local adaptation and potentially identify loci associated with tolerance to abiotic stresses, the polygenic nature of local adaptation (32), suggests that E-GWAS alone may not capture the full extent of genetic variation needed to identify the genetic basis of local adaptation to be advantageous for applications such as plant breeding. There is a similar conceptual and practical advance to be made by moving from GEA and E-GWAS to environmental genome wide selection (E-GS) to estimate the whole-genome adaptive value of an individual to bioclimatic conditions. Making use of environmental variables can extend genomic prediction approaches in meaningful ways that apply to many different species and scenarios.

Crop germplasm collections are a readily accessible resource for landscape genomic studies. However, they often include large numbers of accessions and phenotypic characterization at even a single location may be prohibitive or not possible due to differences in traits such as days to flowering (33,34). This issue motivated the development of the ‘core’ collection concept, which tries to maximize the amount of genetic diversity in the smallest number of accessions (35). The development of core and mini-core collections allows a sufficiently diverse subset of a germplasm collection to be evaluated phenotypically for many traits of interest across different environments, with a cost of missing low frequency beneficial genetic variation. For example, the entire U.S. Department of Agriculture (USDA) barley germplasm collection numbers 33,176 accessions; however, the development of core (n = 2,417) and mini-core (n = 186) collections from this germplasm has allowed practical molecular marker genotyping and phenotyping of different traits (36). Barley landraces are a useful system for exploring this approach because of prior studies of environmental adaptation and the large number of cloned genes associated with adaptation to broad production environments.

Barley (*Hordeum vulgare* L.) is cultivated in an extremely wide geographic and ecological range (from the equator to the inside the Arctic Circle), and has become known as an excellent model for studying and responding to climate change due to its ability to adapt to multiple biotic and abiotic stresses (37,38). In barley, there have been efforts to use GEA and E-GWAS to identify genetic differentiation among wild populations (39), and to find loci associated with abiotic stress tolerance (20,40). Phenotypic recurrent selection and even marker-assisted backcrossing often do not effectively, nor efficiently, transfer quantitative traits into breeding germplasm (41). However, exotic germplasm has been used to explore quantitative (polygenic) traits for centuries, especially in small grains (42). The use of genomic selection provides a theoretical basis for greatly increasing the efficiency of using exotic (unadapted) germplasm as a donor parent to introgress polygenic traits (40). When exploring this strategy in germplasm collections, mini-core collections containing a large proportion of total genetic variation have been shown to be very useful as the initial training population (43). Taking into account the successful application of genomic selection for phenotypic traits, we argue that its use for environmental traits will boost the efficiency in selection of germplasm better adapted to climate change. Barley provides an exemplar source to test environmental genomic selection (EGS) as there are many robust landrace collections (20), and several well-characterized core collections (36). Croplands across the globe are being impacted by climate (1). Previous research has generally shown decreases in yield associated with increases in temperature in barley growing regions (44,45). As a result of these predictions, various adaptation strategies have been suggested, but they have presented limited contributions to breeding populations (46). Further, while genomic selection has become the norm in breeding programs, it has yet to become common in the utilization of germplasm collections, in particular with respect to the use of landscape genomic techniques (47). Here we propose to explore the use of environmental genomic selection (EGS) to identify the best potential parental accessions from the collection of landraces that is maintained by the USDA; thus, improving the speed with which breeding for climate adaptation can occur.

## Results

### Population Structure

The first goal was to recapitulate the previous analysis (20), where four populations had been identified. These relationships were previously established (5,800 SNPs in (20)) and the dataset was divided into central European, Asian, coastal Mediterranean, and East African populations. We identified a similar population structure to that which has been previously identified (20) (using 3,175 SNPs) finding five populations; namely East African (population 1, n=89), Levant/Mediterranean (population 2, n=205), North African/Mediterranean (population 3, n=117), Northern Europe (population 4, n=95) and Asia (population 5, n=278) (**Figure 1; Figure S1**). A comparison of these clusters between the previous study and this study can be found in **Table S2**. These population-clusters recapitulate historic cultivation history, and match well with previous studies.

**Figure 1.**
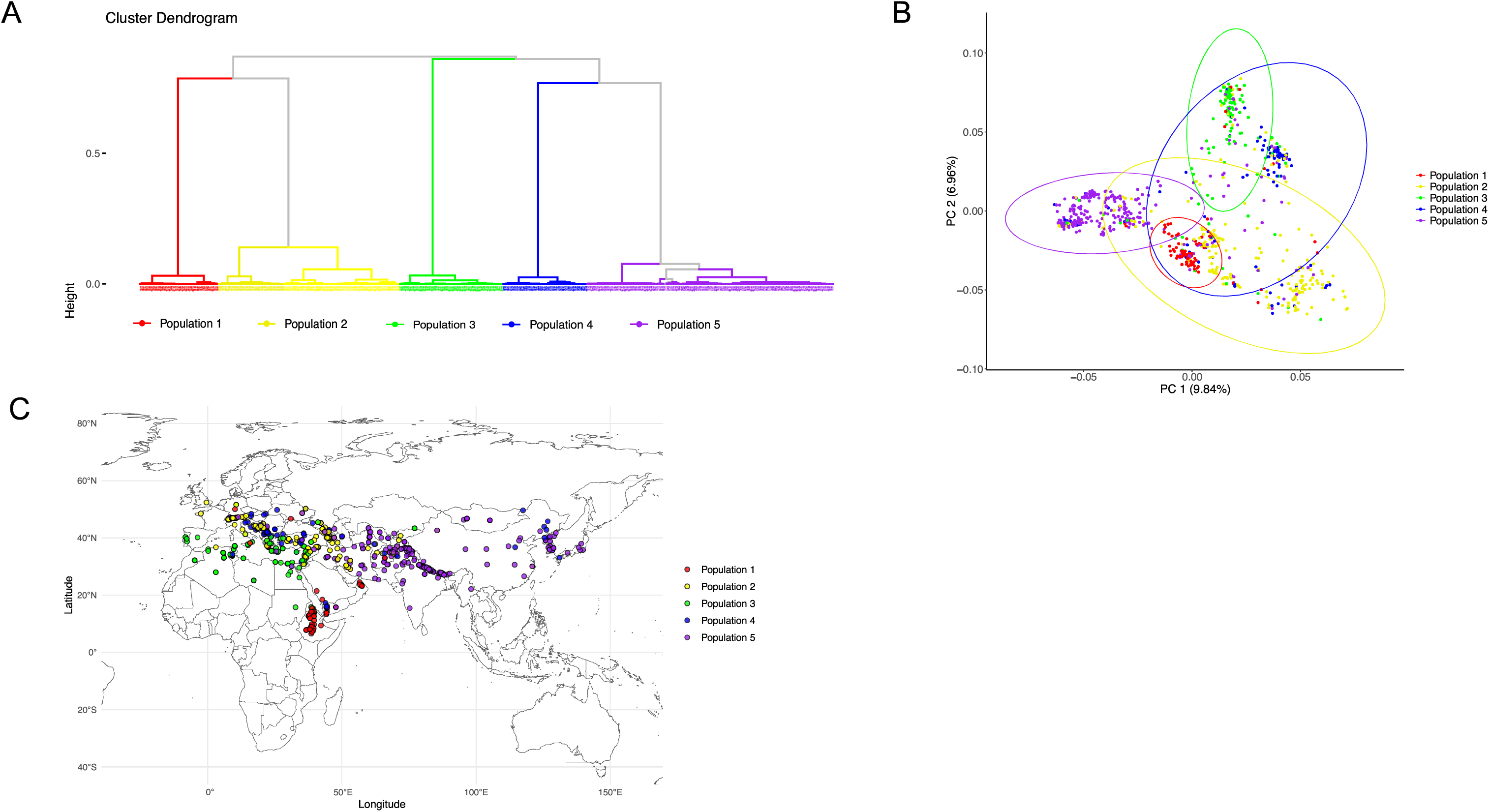
Population structure from Lei et al 2019 dataset **(A)** Hierarchical clustering of 784 landrace barley samples from the Lei et al 2019 dataset based on 3,175 SNPs following an LD prune at 0.2 **(B)** PCA with samples coloured by populations identified in HCPC **(C)** relationship between hierarchical clustering of samples and geographic location.

### Training Population Evaluation and Environmental Genomic Prediction

In this study we explore the utility of core and mini-core collections for genomic selection by using them as the training population for genomic selection calculations. Core and mini-core collections have been previously published for Barley (36). Lines from Munoz-Amatriain *et al* which had the designation of landrace and overlapped with the Lei *et al* publication were used (n=31). Additionally, a de-novo core (n=100) across the 784 lines from Lei *et al* was generated using the corehunter software (48). Prediction accuracy for the mini-core (n=31) and the de-novo core (n=100) was assessed (**Figure 2**). Using 10-fold cross validation with 50 repetitions we assessed four models; rrBLUP, Gaussian Kernel, Exponential Kernal and BayedCPi models. The rrBLUP method was selected as the optimal model for genomic selection. We found good predictive accuracy for the entire collection irrespective of the core size used as the training population (**Figure 2**) with some variables having higher predictive accuracy than others (e.g., bio1, bio3, bio4, bio6, bio11, bio14 and bio17). Exploring the established core collection (n=31) versus the de-novo core collection (n=100), we found little difference in prediction accuracy (**Figure 2; Figure S2; Figure S3**). This shows that as long as a core is developed, prediction accuracies should be high enough for GEAVs to be useful.

**Figure 2.**
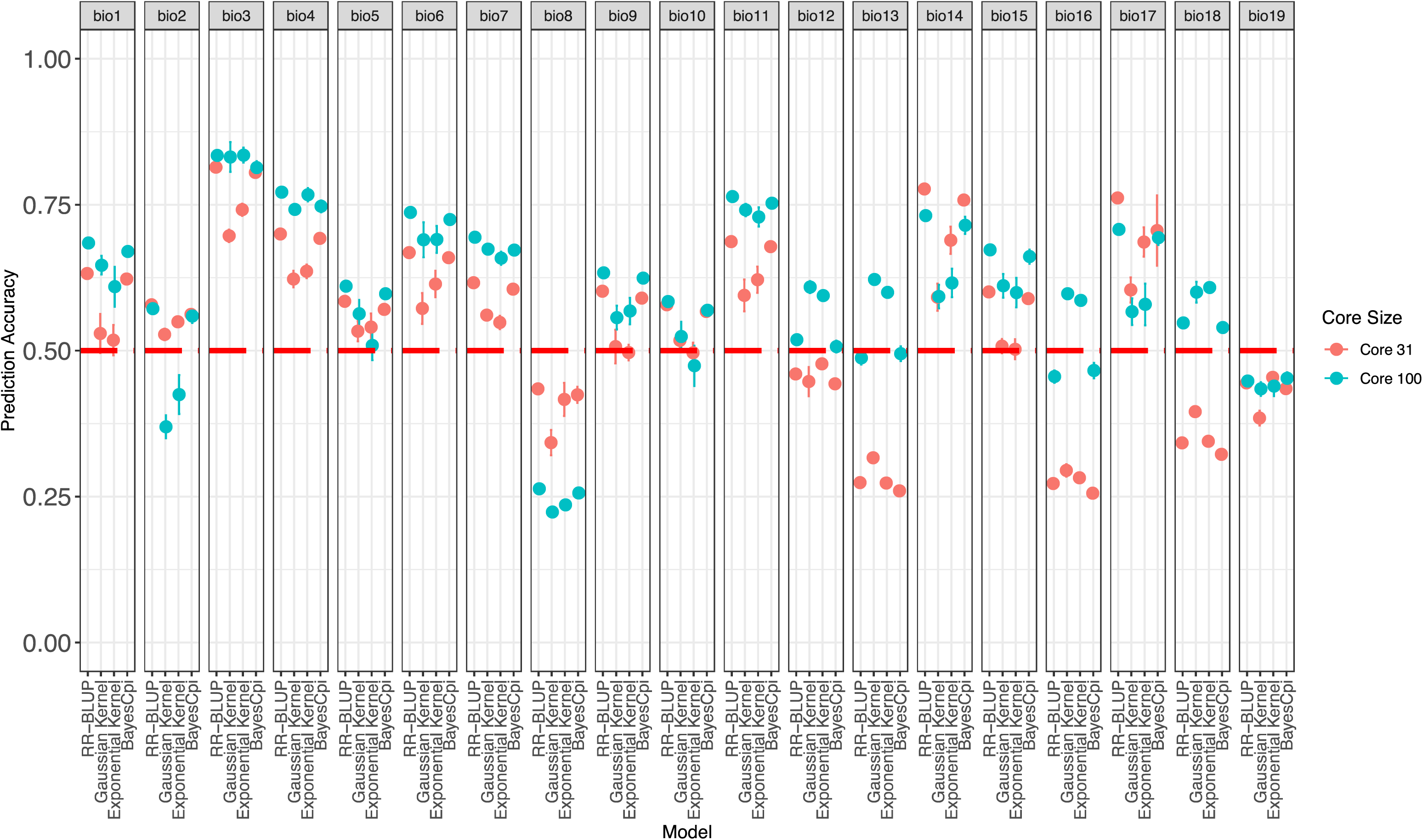
Comparing the n=31 core and the de-novo n=100 core for prediction accuracies across models.

Genomic prediction using rrBLUP was performed on both of the core collections on the variables which had higher prediction accuracy (bio1, bio3, bio4, bio6, bio11, bio14 and bio17) and GEAV values were assessed for each line (**Table S3; Table S4**) and population (**Figure 3A**; **Figure 3B**). We found good predictive accuracy for the entire collection (**Figure S2**) despite the low number of individuals (n=31) used as the training population. Some variables had higher predictive accuracy than others and there was limited overlap for multiple environmental variables. For some variables, there were wider distributions of GEAV values (e.g., bio4 - temperature seasonality **Figure 3**), which indicates that for some environmental stressors, landrace accessions have more potential to adapt, but there is not a straightforward interpretation in all cases. It was clear that there were different variables that had higher predictive accuracy in different populations (**Figure 2; Figure S3**). However, when exploring the established core collection versus the de-novo core collection, we found little difference in prediction accuracy (**Figure 2; Figure S2; Figure S3**). This shows that as long as a core is developed, prediction accuracies should be high enough for GEAVs to be useful. The distinct evolutionary histories of the populations may mean that to fully take advantage of GEAV, decoupling the population structure may provide a more accurate assessment of which accessions may be the best line. For example, East Africa (population 1) has poor GEAV values for bio 4 (temperature seasonality), but Asia (population 5) has the highest mean GEAVs for this climate variable (**Figure 3**).

**Figure 3.**
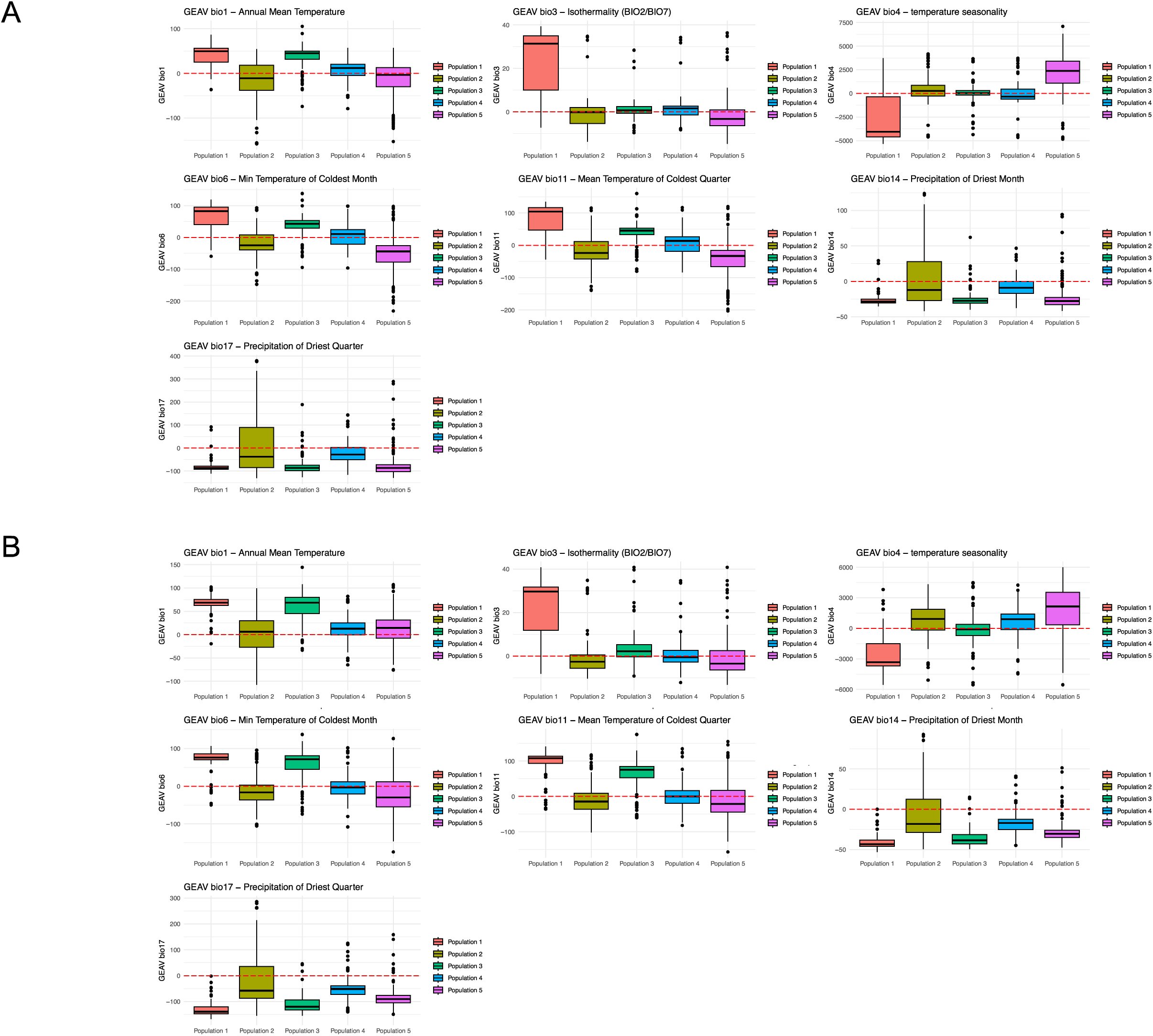
Population-level GEAV values for climate variables with the highest prediction accuracy. GEAV values are grouped and colored by hierarchical clustering within the dataset (k=5) **(A)** for training set n=31 **(B)** for training set n=100.

When exploring the overlaps in accessions in the top 5% of GEAVs for temperature related climate variables (bio1, 4, 6 and 11) specialists were identified in non-overlapping regions (bio1 (n=15), bio11 (n=1) and bio4 (n=40) (**Figure S4A**). Specifically, we detected a large overlap among accessions with high GEAVs for bio1 - mean annual temperature, bio 6 - minimum temperature of the coldest month, and bio11 - mean temperature in the coldest quarter, but poor overlap with bio4 - temperature seasonality (**Figure S4A**). Similarly exploring the lines that had high GEAVs for precipitation namely bio14 and bio17, a complete overlap in the accessions was seen in the top 5% of GEAV values (**Figure S4B**), with 32/40 coming from Levant/Mediterranean population (population 2), 1/40 from North African/Mediterranean (population 3), 3/40 from Northern Europe (population 4) and 4/40 lines from East Asia (population 5).

### Environmental Variation and GEAV association

The populations defined by genetic assignment differ in terms of which environmental variables are most strongly associated with population-level variance. When looking at all the populations together, we observed that distinct variables were more associated with each population (**Figure 4A** ; **Figure S5**). For example, bio4 - temperature seasonality was associated with the Asian population (population 5) (**Figure S5E**), while bio17 - Precipitation of Driest Quarter was more associated with the Levant/Mediterranean population (population 2) (**Figure S5B**). By plotting GEAV values for environmental variables at each line’s geographic origin, patterns in GEAV distribution become apparent. For example, for bio3 - isothermality more southern latitudes and more specifically lines from population 1 (East African population) have high GEAVs for this trait (**Figure 4C**; **Figure 3A/B**). In contrast, when breeding for temperature seasonality – bio4 (**Figure 4D**) lines in more northern latitudes and more specifically population 5 have high GEAVs for this trait (**Figure 3A/B**).

**Figure 4.**
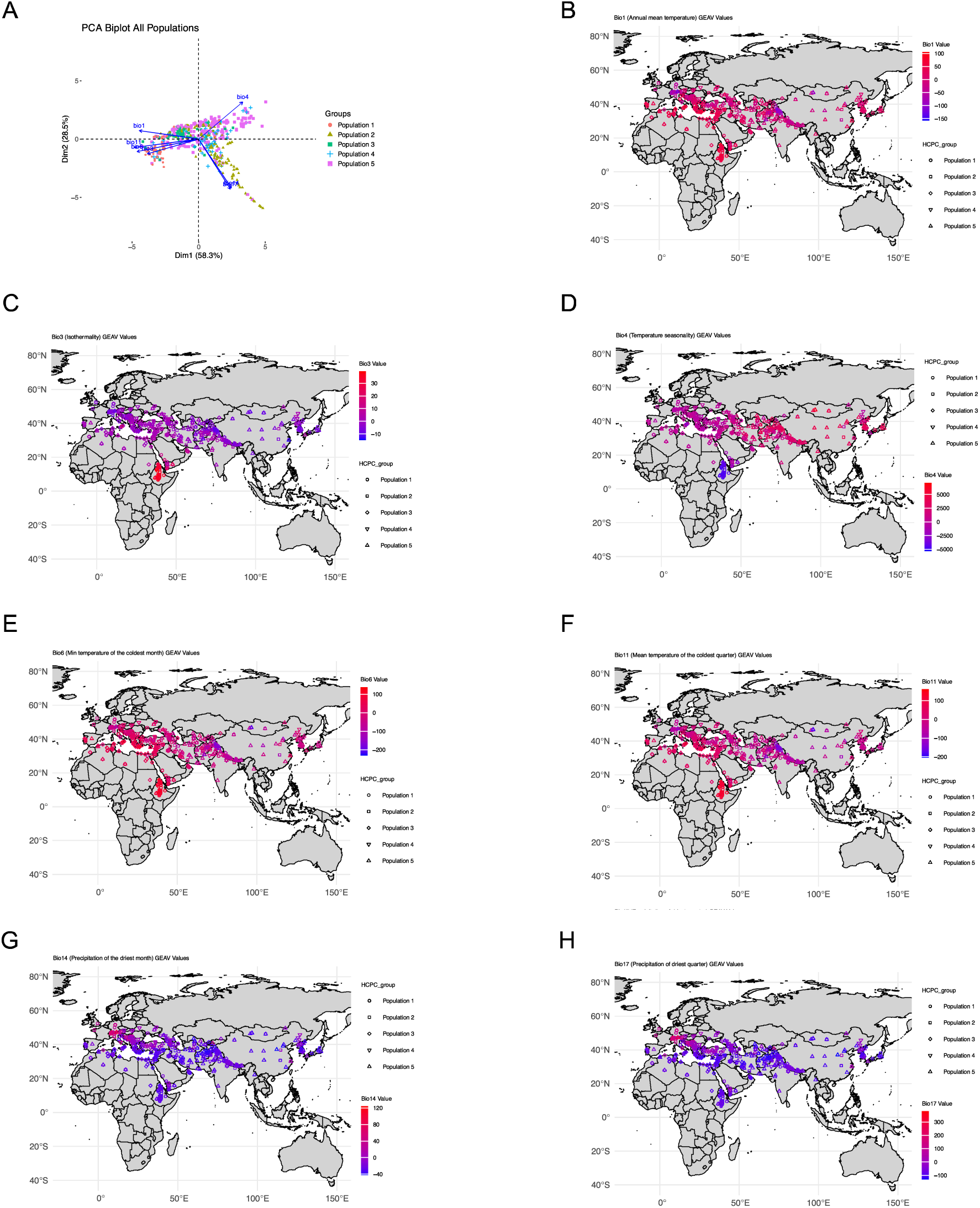
Relationship between environmental variables and geographic distribution of GEAV values by population for each climate variable **(A)** Relationship between environmental variables and lines associated with each population **(B)** Geographic distribution of GEAV values for Bio 1 - Annual mean temperature **(C)** Geographic distribution of GEAV values f or Bio 3 - Isothermality **(D)** Geographic distribution of GEAV values for Bio 4 - Temperature seasonality **(E)** Geographic distribution of GEAV values for Bio 6 - Min temperature of the coldest month **(F)** Geographic distribution of GEAV values for Bio 11 - Mean temperature of the coldest month **(G)** Geographic distribution of GEAV values for Bio 14 - Precipitation of the driest month **(H)** Geographic distribution of GEAV values for Bio 17 - Precipitation of the driest quarter.

### Leveraging population distribution models

The geographic coordinates from each individual within each population were used to create population distribution maps (PDM), similar to previous work in common bean (49) and created a new response factor (suitability score for each sampling location). The different environmental characteristics of each PDM may be driving genetic architecture and population divergence and thus could be explored for favorable alleles in parent selection for breeding for potential future environments. There were clear differences in optimal areas for each population identified (**Figure 5; Figures S6-10**). For example, the East African population (**Figure 5B; Figure S6**) showed a narrower range than populations from Northern Europe (**Figure 5E; Figure S9**) and Asia (**Figure 5F; Figure S10**). Sources of local adaptation can be found in each of the populations and suitability for different environmental variables (**Figure S6-10**). Each pixel in the suitability model shows the habitat conditions of the geography which greatly impacts the ability of a population to grow in that region, typically if a value is above 0.2 plants will be able to grow with agricultural intervention. We further explored the GEAV values for SDM suitability scores (**Figure S11**). Prediction accuracies were highest for the specific population the SDMs originated from, with the exception of population 4 (**Figure S11A**). Additionally, population distribution models can be utilized to identify key parental lines for breeding programs aimed at adapting to specific environmental conditions. By intersecting distribution ranges with lines that fall within these ranges (**Figure 5B-F; Table S9-S13 (core n=31) and Table S14-18 (core n-100)**), breeders can select genetic backgrounds that are optimal for their environment or target environment. Selecting lines with high GEAVs for the trait of interest in these ranges can minimize linkage drag and expedite introgression of favorable alleles (**Table S9-S18**).

**Figure 5.**
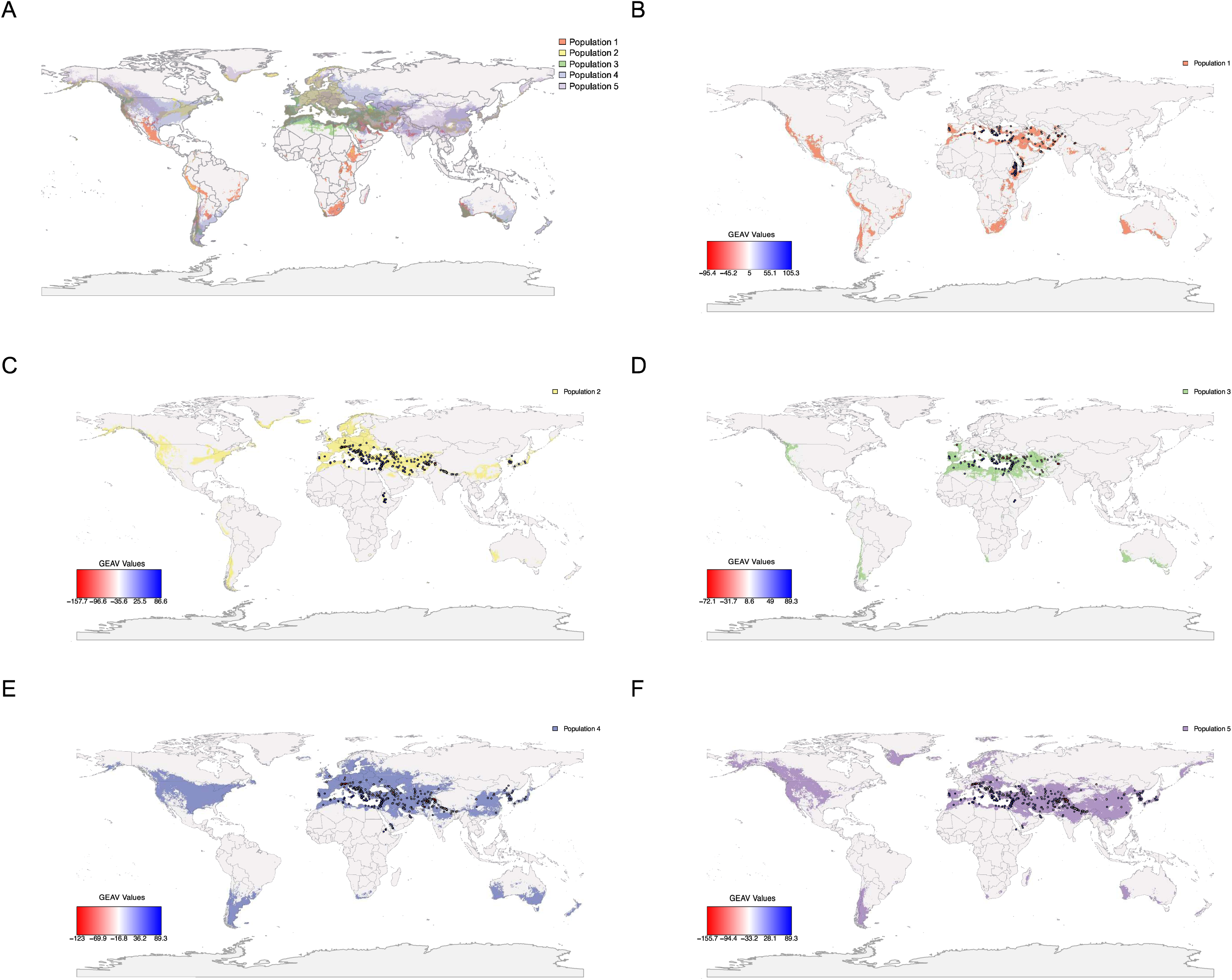
Intersection of Species Distribution Modelling and population level GEAVs **(A)** Overlaps in species distribution for all five populations **(B)** Population 1 - East African population specific species distribution with range specific lines for bio1 **(C)** Population 2 - Levant/Mediterranean population specific species distribution with range specific lines for bio1 **(D)** Population 3 - North African/Mediterran ean population specific species distribution with range specific lines for bio1 **(E)** Population 4 - Northern Europe population specific species distribution with range specific lines for bio1 **(F)** Population 5 - Asian population specific species distribution with range specific lines for bio1.

### Chromosomal patterns of adaptation across populations

We examined chromosomal patterns across the five populations. When exploring individual marker scores there were distinctly different patterns across the populations (**Figure 6; Figure S12; Figure S13**). For example, the pattern on chromosome one is distinctly different in the Levant/Mediterranean population than the other populations, while the East African population shows a different pattern on chromosome 2 (**Figure 6**). This shows that despite many of the response factors being highly quantitative there are patterns that are genetic background specific that can lead to more local adaptation. Despite these different patterns there were often common SNPs that had consistently large effects for specific traits (**Table S5; Figure S14; Supplemental File 1**). While these markers had larger predicted effects on the phenotype (e.g., sometimes being related to a change in temperature of ∼0.1°C) these SNPs did not always follow the geographic distributions of the populations, often the SNPs would have patterns where the beneficial allele would be present in many populations (**Figure S14**). Marker effects were not distributed evenly across chromosomes or among populations (**Figure S12; Figure S13**). There seem to be many private alleles within populations that have an adaptive effect (**Figure S13**). These variants are not evenly distributed by trait, suggesting that some populations apparently have more tolerance than others for specific environmental stressors.

**Figure 6.**
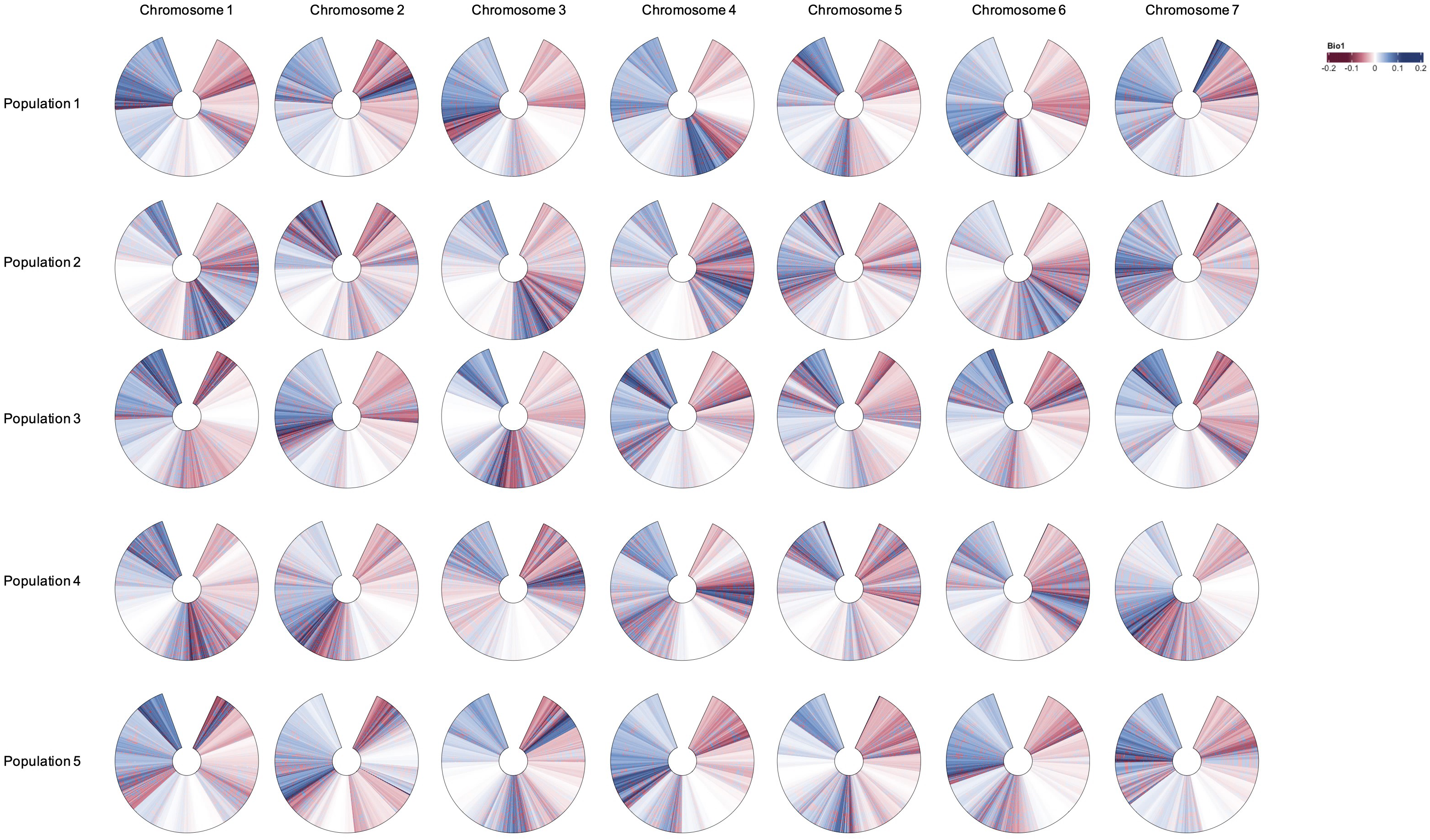
Population-level chromosome painting for bio1 for East African (Population 1, n=89), Levant/Mediterranean (Population 2, n=205), North African/Mediterranean (Population 3, n=117), Northern Europe (Population 4, n=95) and Asia (Population 5, n=278).

## Discussion

### Adapting to Climate Change

Using a large collection of publicly available landraces of barley that contained georeferences and were genotyped with a common SNP panel, we explored which individuals may be the best parents for use in breeding programs targeting environmental stress adaptation. Genomic prediction was conducted using a subset of the barley core collection that was represented in the georeferenced accessions as the training population. Further, to try and better understand the potential for different climate niches, population specific distribution models were created to explore if parental material from population specific niches would be more useful. In general, we found specific accessions that had high GEAVs across a wide range of variables (temperature, precipitation, climate niche), indicating there are some parents that have much higher potential. Next, we found that there were often population specific accessions that had high GEAVs, suggesting that linkage drag could be minimized by selected parents that are more closely related to your breeding material, while still gaining most of the abiotic stress benefits. Finally, there were often population specific patterns of variation, indicating that there is potential for combining different genetic architectures to create more resilient plants, with individuals with high GEAVs likely to have different polygenic adaptations. The approach explored here is broadly applicable to a range of problems, including selection of trees for plantations, conservation efforts (e.g., using genomic prediction instead of genetic offsets) and other crops. With the decreasing cost of generating sequencing and improvements in the ability to get geographically relevant information, the analytical approach here can be applied to many systems. It is also important to understand the distribution of GEAVs as recent work has shown that there is increased genetic load at range edges (50) this has implications for where individuals for future breeding work should be selected from.

### Exploration of Previously identified Candidate Genes

In previous work, known genes were identified with large impacts on local adaptation (20,51). In particular, there were often allelic series identified across populations where in different environments different alleles were favored (51). When exploring the GEAVs specific genes had a disproportionate impact on the total variance explained. Although some of these genes were previously identified through GEA analysis they were in the extreme ends of the distribution for marker effects (**Table S6; Figure S15),** and explained a small amount of variance. This suggests that climate change adaptation will be largely polygenic and that individual genes will not be strong enough for these environmental responses. This suggests that a marker assisted backcrossing approach to create resilience will be less successful than a genomic prediction approach. Using the model where we treated candidate genes from previous GEA analysis as fixed effects the model fit was improved, but there was not an improvement in prediction accuracy using the either core collection when fixing cold tolerance (**Figure S16A**) and flowering time (**Figure S16B**) related genes. These candidate genes had different impacts on prediction accuracy for different traits, which is expected based on the relationship between a particular gene and the environmental response of interest. Unlike previous work in rice (52) and wheat (53), but similar to previous simulation work (54) we found that using fixed effects did not improve prediction accuracy (**Figure S16**). This could be due to the highly quantitative nature of the abiotic stress tolerance, or it could be that due to overfitting the model by including fixed effects that are not related to the specific abiotic stress. In either case, it seems like the best approach will likely be to use a genomic prediction method rather than a marker assisted backcrossing method for adaptation to climate change.

### Maximizing the efficiency of selection to incorporate exotic germplasm for adaptation to climate change: How much progress can be made in creating locally adapted cultivars?

Plant breeding takes a long time, depending on the plant system it can take between 8-20 years to release a new cultivar (55). Breeding programs have long appreciated the utility of collecting and storing genetically diverse accessions of crops and their wild relatives to maintain the genetic variation essential to breeding progress (56,57). Germplasm collections of the wild relatives of crops have had great value to agriculture by providing, for example, new alleles for disease resistance or crop quality (58,59); however, they have been underutilized with respect to climate change (15,60,61). While many studies have examined approaches to best exploit germplasm collections in plant breeding programs (43,62), optimizing the selection of accessions remains a challenge. Currently, breeding programs are not releasing cultivars fast enough to keep up with predictions for climate change (63). This implies that the evolutionary relationships between populations can perhaps provide insight into adaptation in modern cultivars to future climate change, due to understanding how both genetic background and loci of large effect interact in populations that were the founders of proximate modern cultivars. It is clear that targeted decisions can greatly increase the speed of breeding. New developments in high-throughput phenomics (64), high-throughput genotyping (65) and speed breeding (66) may provide ways to rapidly introduce novel abiotic stress tolerance genes. Further, combining environmental genomic selection with other metrics of local adaptation, for example home field advantage (67), i.e., combining locally adapted with environmentally adapted material, may be a pathway to more rapidly develop cultivars for specific geographies. EGS is an extension of GEA analysis to make diverse collections more available to plant breeding programs. While classic GEA analysis provides important information about specific alleles that are putatively adaptive, it does not provide direct information about parental line performance or value as a parent (7). Traditional genomic selection has been employed to decrease cycle time (68), extending this approach by exploiting local adaptation should enhance breeding for climate change. EGS should provide better parental selection because instead of focusing on loci of large effect that may be in an unadapted genetic background, it can incorporate more quantitative information. Also, this approach still allows for a mechanistic understanding while pushing forward populations within active programs, in effect it allows for both a retrospective and prospective exploration at the same time.

### How do you update your training population in the local breeding program?

Historically heat and drought stress have been very difficult to phenotype (69). Recent advances in controlled environment agriculture and phenomics have increased the measurement precision (70), which combined with the advent of artificial intelligence in breeding (71) are expected to advance breeding for climate change. These two phenotypes are projected to be some of the most important traits under climate change. The difficulty of phenotyping these traits make them excellent candidates for GEA and EGS, however, once parents are selected and incorporated into breeding programs the initial training population will no longer be the most appropriate (72). The GEAV approach can be explored for any particular climate response (**Figure 4; Table S3-4**). For example, while drought is a problem in North America, in northern Europe waterlogging/flooding is likely to be a larger problem. Here for the precipitation related climate variables examined (bio14 and bio17) we can see that the highest GEAVs for these traits occur in regions that have high precipitation, even in dry parts of the year (**Figure 4G**; **Figure 4H)**. Given the difficulty in phenotyping abiotic stress, ensuring that continued progress can be made in the next cycle, it is important to update the training population. The first step in doing this will be making sure that your elite parents have been phenotyped for abiotic stress tolerance in the normal way it is assessed in the breeding program. Further, it will be important to test multiple training populations to optimize resource allocation for a specific breeding program. A major next step will be to understand if you can continue to select for abiotic adaptation after more than one generation of crossing or if this method is best suited for parental selection.

## Caveats

It is important to note that in this study there was limited marker coverage, which may impact the overall GEAVs. For example, the IPK germplasm collection has been genotyped with GBS having much higher marker resolution (73). While we speculate on the best way to incorporate this exotic germplasm into breeding programs it will be important to conduct both simulation and empirical studies to make sure the rate of gain per cycle is similar to marker assisted selection. The GEAV calculation is based on historic mean values for climate variables from 1970-2000 (74). Having more accurate climatic data with or making use of the entire time-series would lead to better results.

## Conclusion

Barley is grown from the tropics to the Arctic Circle. Despite this large range there are clear populations which differ greatly in their predicted distributions. These different populations have different genetic values (GEAVs) for breeding for climate change. It is clear that large scale genotyping of landrace material followed by genetic characterization with environmental genomic selection can identify promising parents and reduce the time required for the breeding process. Here we have used publicly available data (genotypes, with georeferences for accessions and worldwide climate data) and identify landraces that have shown polygenic adaptation to climate niches and specific environmental variables and likely host beneficial alleles for introgression when breeding for target environments.

## Materials and Methods

### Data Sources

#### Core Collection/Genotype

The USDA barley core collection comprises 2,417 accessions (36). Based on 9K Illumina Infinium iSelect Custom Genotyping BeadChip (75), a set of 1,860 non-redundant samples were retained, identified as the iCore. These accessions originated from 94 countries and included 815 landraces. From the iCore collection, Muñoz-Amatriaín *et al*. further developed a mini-core collection comprising 186 accessions, 31 of which were landrace samples, which are represented here as the n=31 core. Genotypes reported here derive from automated genotype calling implemented in the software Alchemy (76). SNP calls with posterior probability >0.95 were retained, while calls below the threshold were treated as missing data (42). The VCF file used for analysis here was reported in Lei *et al*. (20) using SNP physical positions in the Morex_v2 assembly (77). Lei *et al*. (20) selected 803 landrace accessions from the iCore and following quality filtering and the exclusion of accessions lacking distinct locality information, they identified a final set of 784 georeferenced landrace accessions, which was selected as the genetic material in this study.

#### De-novo core collection methods

The genotypic data from the Lei et al., (20) was provided in Supplemental_dataset_1.vcf. This VCF was converted into a genlight object using the ‘vcfR’ (78) and ‘adegenet’ packages (79). A distance matrix was calculated using the ‘poppr’ package (80). Hierarchical clustering was performed on the distance matrix using the Ward method, and the dataset was clone-corrected to account for potential duplicates. To show this method would be applicable to species where no core was developed, we also developed a de-novo core of the 784 lines. This was generated using the ‘corehunter’ package (81) where 100 lines (core n=100) were sampled from the precomputed distance matrix.

#### Population Structure and Environmental Genomic Selection

The above dataset was examined for population structure using ‘SNPRelate’ (82). For the Principal Component Analysis (PCA) and Hierarchical Clustering of Principal Components (HCPC) only bi-allelic SNPs further filtered for linkage disequilibrium (0.2) (3,175 SNPs) were used. Genomic prediction was performed using 6,068 SNPs to predict bioclimatic and biophysical variables to generate a genomic estimated adaptive value (GEAV) for each accession for a given trait (conceptualized as the genetic value for a specific environmental context) (**Table S3; Table S4**). In previous work these have been characterized as genomic estimated adaptive values (GEAVs - (47). Four genomic prediction methods were examined: i) RR-BLUP, ii) G-BLUP with an exponential kernel, iii) G-BLUP with a Gaussian kernel, and iv) BayesCπ. R packages ‘rrBLUP’ (83) and ‘hibayes’ (84) were used for the analysis (**Figure 2**; **Figure S2**). The training population (core n=31) consisted of 31 georeferenced accessions, representing the overlap between the mini-core collection identified by (36) and the 784 landrace samples used in (20). The remaining 753 landraces from the Lei et al. (20) dataset was used as the validation set. Prediction accuracy was based on Pearson correlation (r(PGE,y)) between the predicted genotypic effects and the observed environmental variable with 10-fold cross-validation. Environmental traits which were ascribed a prediction accuracy over 50% for the majority of methods were examined in more depth (**Figure S2A**). For the de-novo core (core n=100) prediction accuracy was examined similarly with the remaining 684 landrace lines used as the validation set (**Figure S2B**). When setting famous genes of interest as fixed effects (n=22) in the prediction and genomic selection models, highly correlated SNPs were removed due to multicollinearity using the R package ‘caret’ (85). This resulted in 14 SNPs which were set as fixed effects. Prediction accuracy for the rrBLUP model with these fixed effect SNPs was calculated for both cores (n=31 and n=100) (**Figure S16**) as well as genomic selection and GEAVs (**Table S7; Table S8**). Rank changes across the environmental variables were also examined (**Figure S17**).

#### Environmental Data

Occurrence data from Munoz-Amatriain *et al.* (36) were separated into populations based on the genetic assignment analysis (see above). This led to 89 individuals in population 1, 205 individuals in population 2, 117 individuals in population 3, 95 in population 4, and 278 in population 5. These occurrence points were used to query the WorldClim 2.1 climate data (all 19 bioclim variables for temperature and precipitation, **Table S1**). Data was downloaded at the highest available spatial resolution of 30 seconds (∼1 km) (https://www.worldclim.org - (74)). These bioclimatic data were used to create species distribution models (SDM) using the software Maxent (Version 3.4.4 - (86)) in RStudio (Version 2022.2.0.443 - (87)). Map overlays were created using the ‘raster’, ‘rworldmap’, ‘ggplot2’, ‘sf’ and ‘mapdata’ packages in RStudio. Suitability maps were overlaid for the present day (1970–2000), with a suitability cutoff score of 0.2. Acceptable suitability is defined as 0.2 for cultivated regions (88) and 0.4 for natural areas (89). Model quality was explored using the area under the curve (AUC) and the standard deviation of the AUC across replicates (SDAUC). A good model requires an AUC ≥ 0.7 and SDAUC < 0.15. The final SDM suitability value was then used as a response factor for environmental genomic selection.

## Supporting information

Tabld S5

## Data and Code availability

Data are available at https://github.com/ahmccormick/Barley_EGS

