## Supplemental Figures for "Back to the Future: Environmental genomic selection to take advantage of polygenic local adaptation"

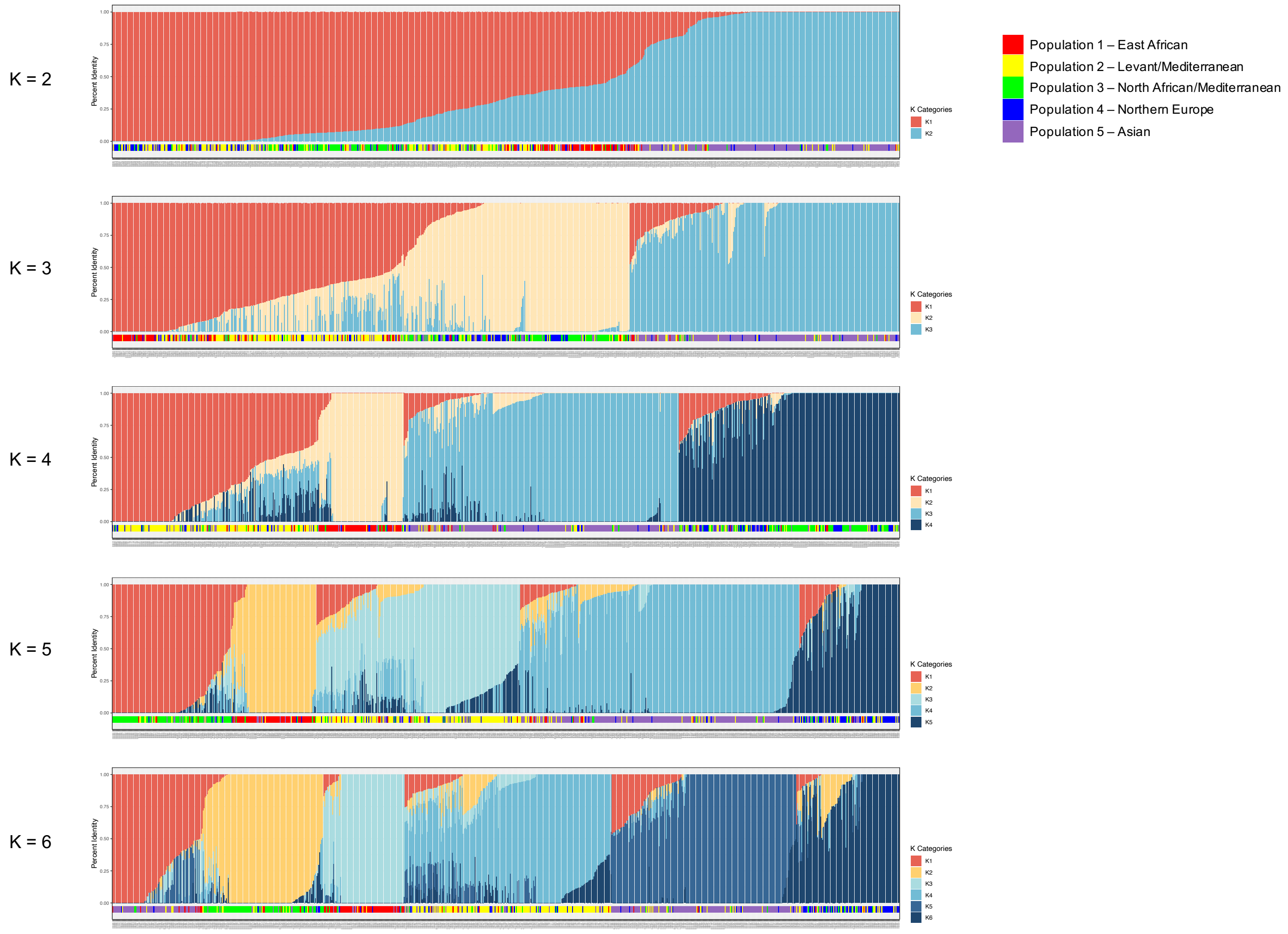


***Figure S1.*** *Visualization of population structure and admixture from 6,068 SNPs using the fastSTRUCTURE software (k=2-6).*

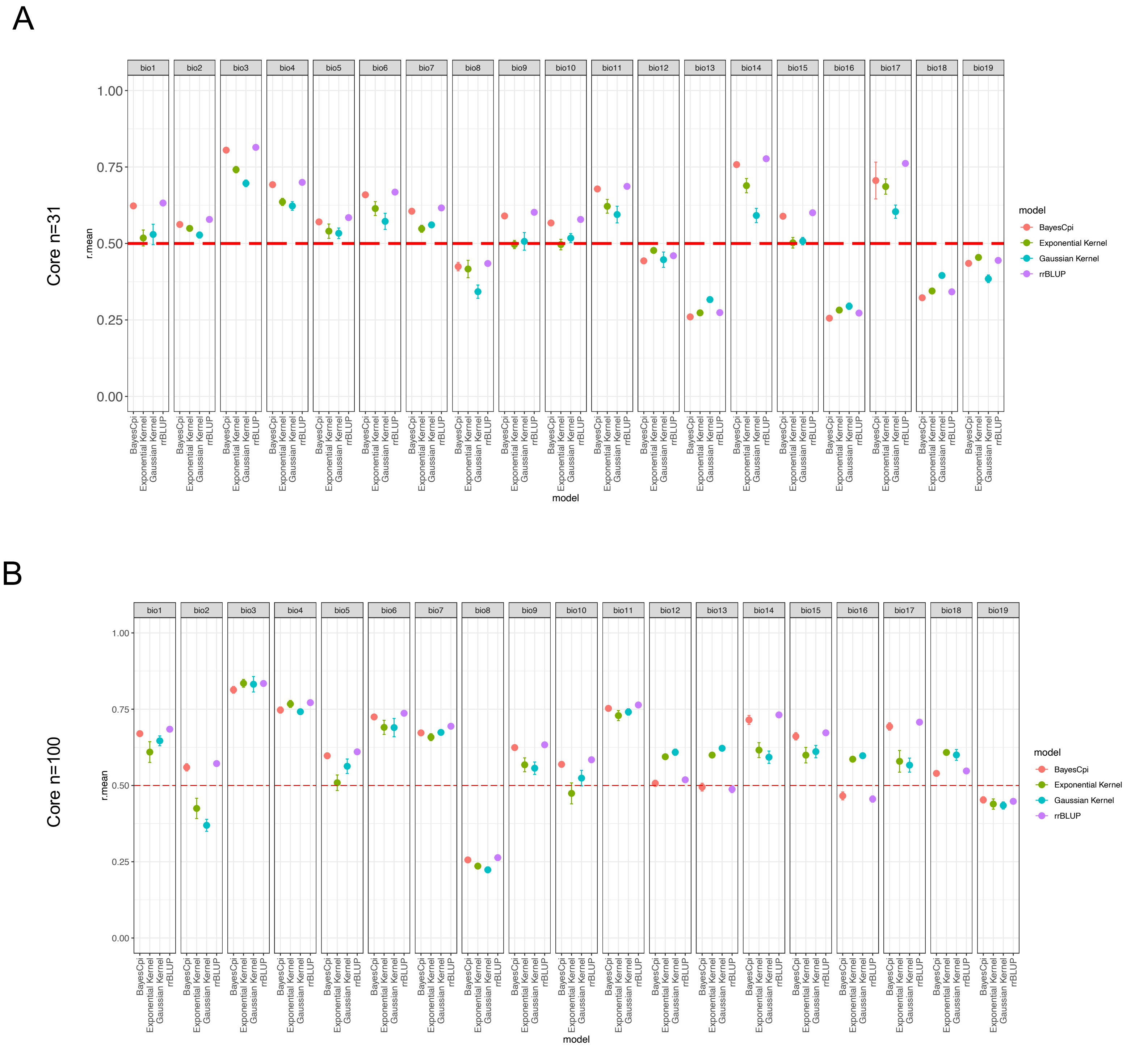


***Figure S2***. *Prediction accuracy for WorldClim climate variables using four genomic prediction methods. rrBLUP, G-BLUP with an exponential kernel, G-BLUP with a Gaussian Kernel and BayesCpi at k-fold = 10* ***(A)*** *A training set of 31 lines was used as this was the overlap between the minicore (Munoz et al., 2014) and the lines used here from Lei et al., 2019) leaving a test set = 753 as compared to* ***(B)*** *A training set of 100 lines was identified from core hunter, with a test set of 684 lines.*

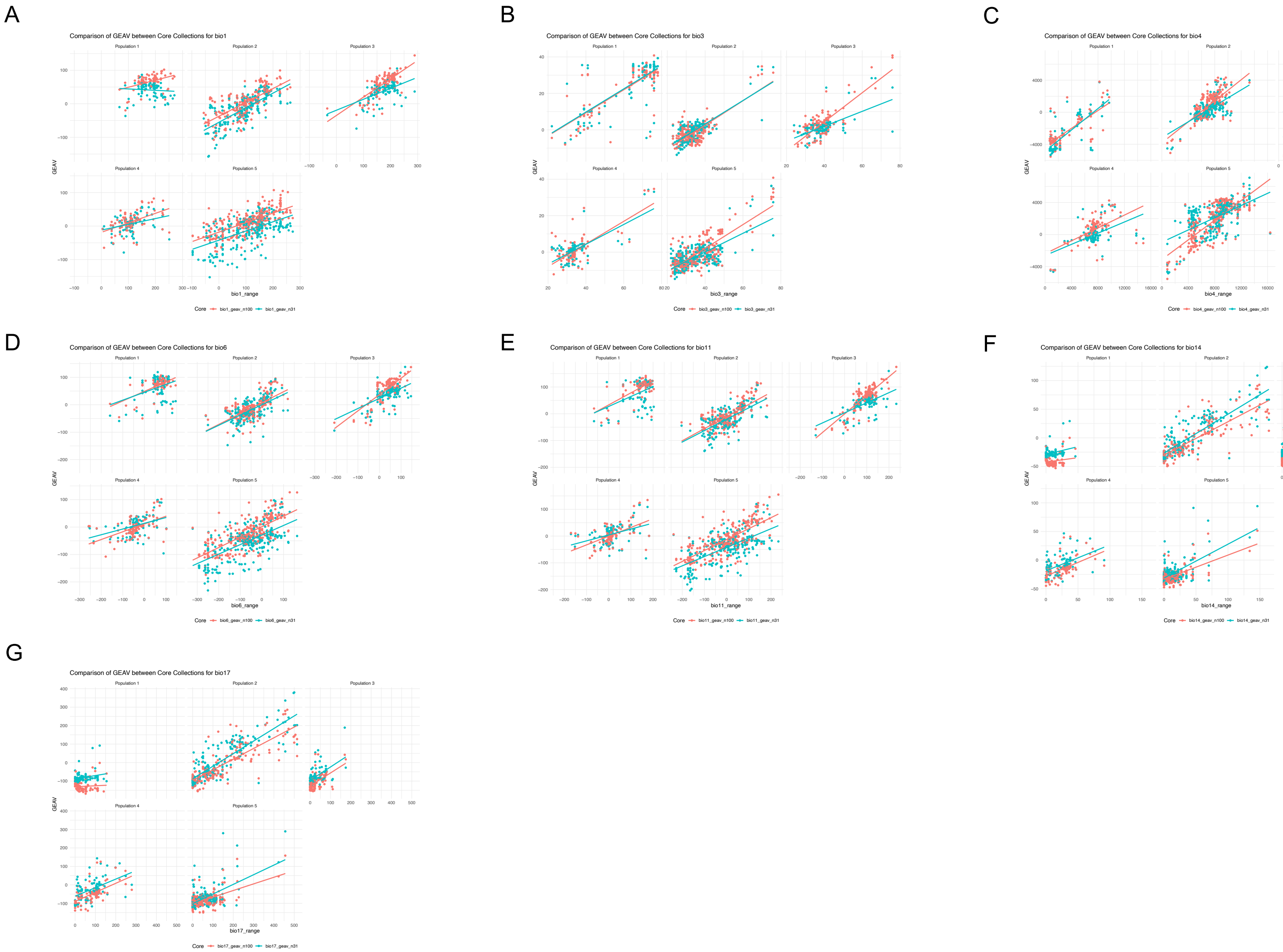


***Figure S3.*** *Examining the n=31 core and de-novo n=100 core for GEAV consistencies per population for bioclimatic variables **(A)** Bio1 (Annual Mean Temperature) **(B)** Bio 3 (Isothermality) **(C)** Bio 4 (Temperature Seasonality) **(D)** Bio6 (Min Temperature of Coldest Month) **(E)** Bio11 (Mean Temperature of Coldest Quarter) **(F)** Bio14 (Precipitation of Driest Month) **(G)** Bio17 (Precipitation of Driest Quarter).*

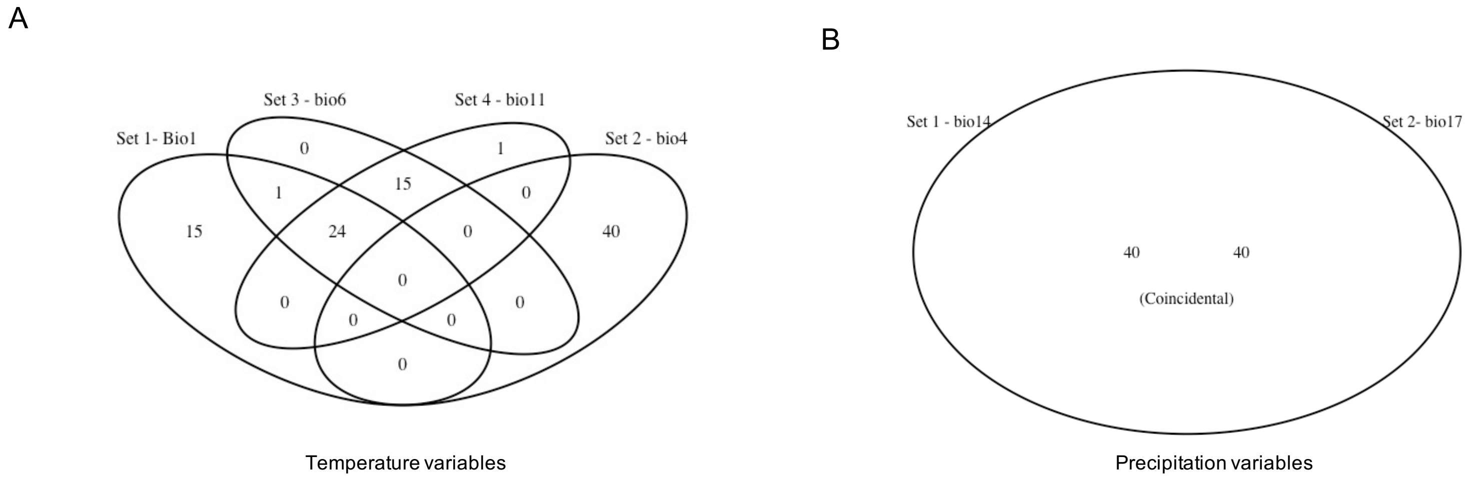


***Figure S4.*** *Overlap of the top 5% of lines with the highest GEAV values within each climate variable* ***(A)*** *temperature variables (bio 1, 4, 6, 11) and* ***(B)*** *precipitation variables (bio 14,17). Specialists for temperature-related variables were identified in non-overlapping regions (bio 1 (n=15), bio11 (n=1) and bio4 (n=40). Precipitation overlaps show complete congruence for lines.*

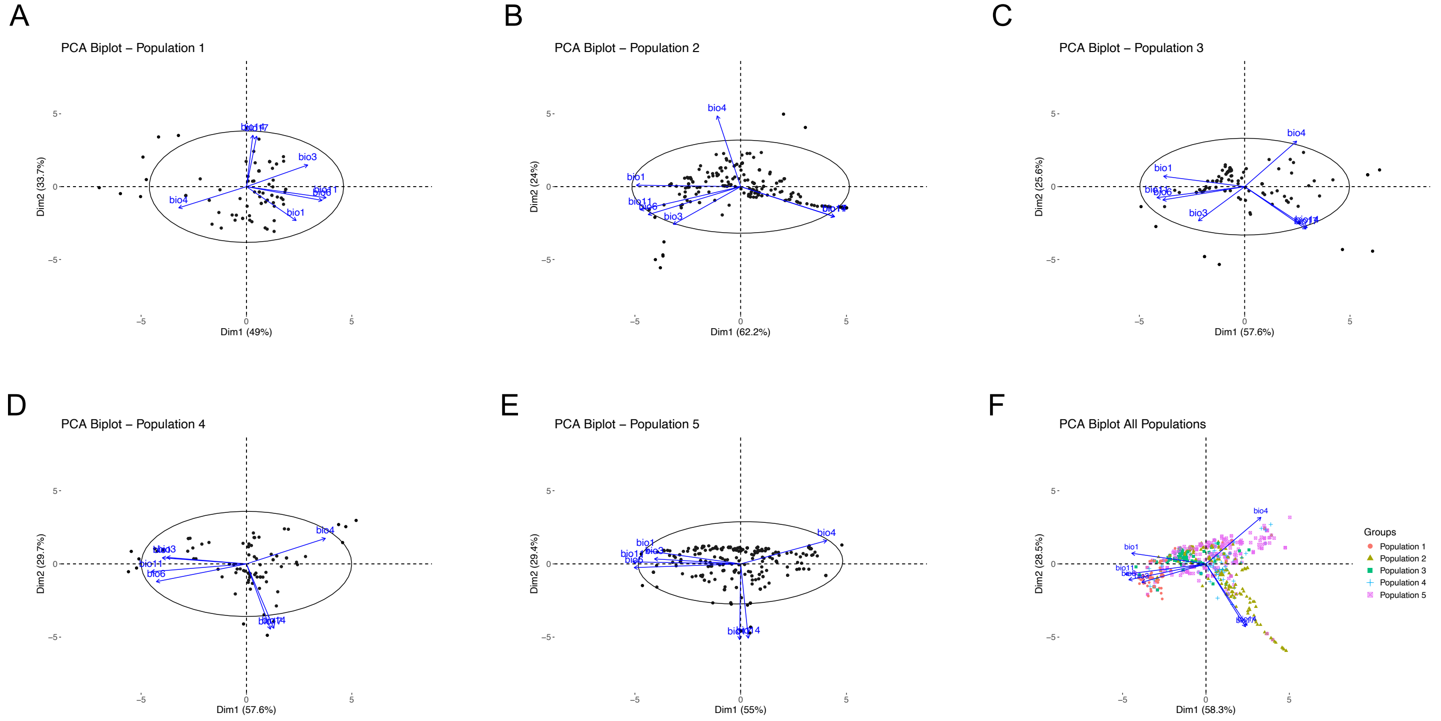


**Figure S5.** *Relationship between environmental variables and lines associated with each population* ***(A)*** *Population 1 - East African population* ***(B)*** *Population 2 - Levant/Mediterranean population* ***(C)*** *Population 3 - North African/Mediterranean* ***(D)*** *Population 4 - Northern Europe* ***(E)*** *Population 5 - Asian population* ***(F)*** *Relationship between all environmental variables and 784 lines in the dataset.*

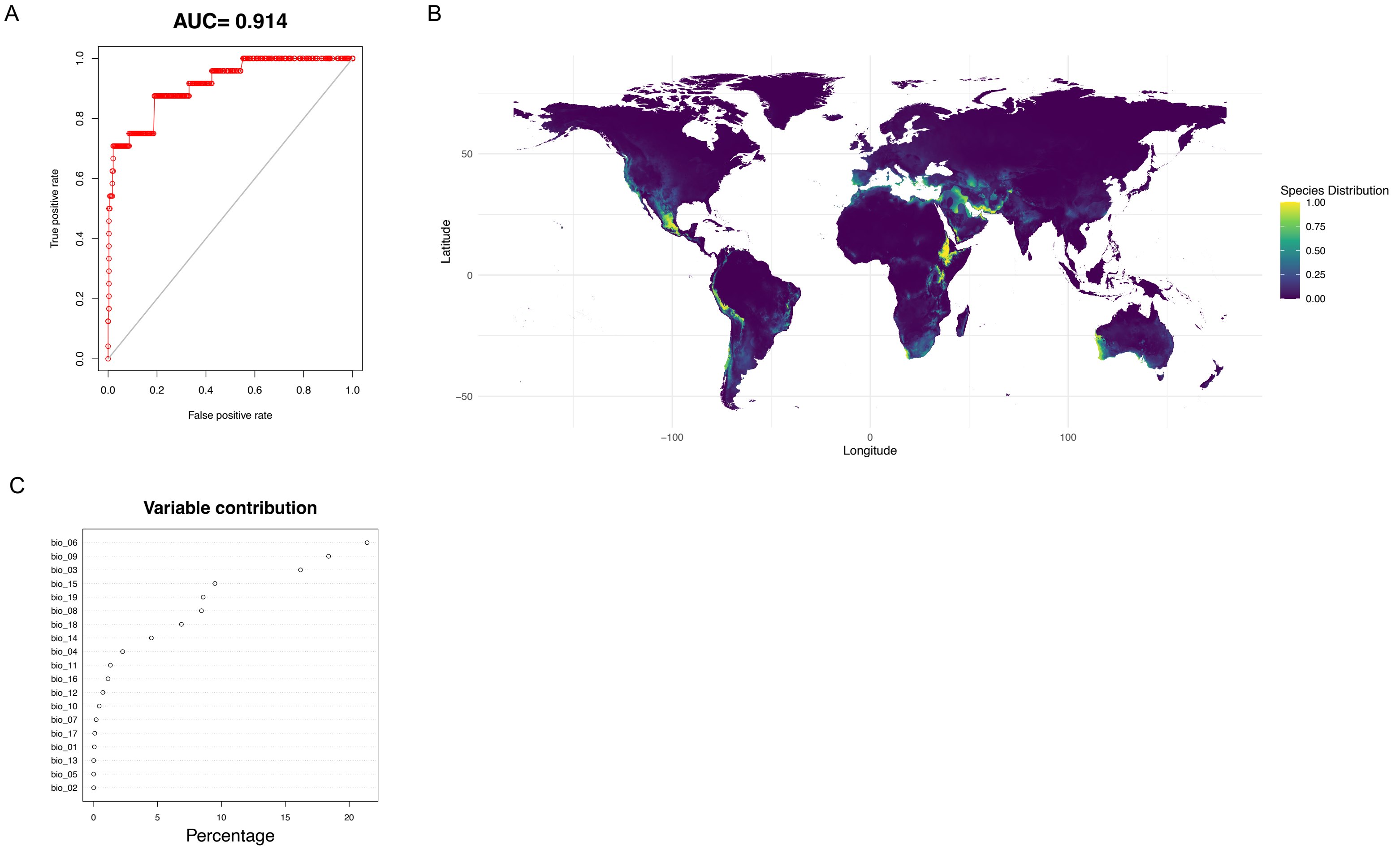


***Figure S6.*** *Species distribution modelling results for population 1* ***(A)*** *Area under the curve (AUC) graph* ***(B)*** *Species distribution map and* ***(C)*** *Variable contribution graph for the WorldClim bioclimatic variables examined.*

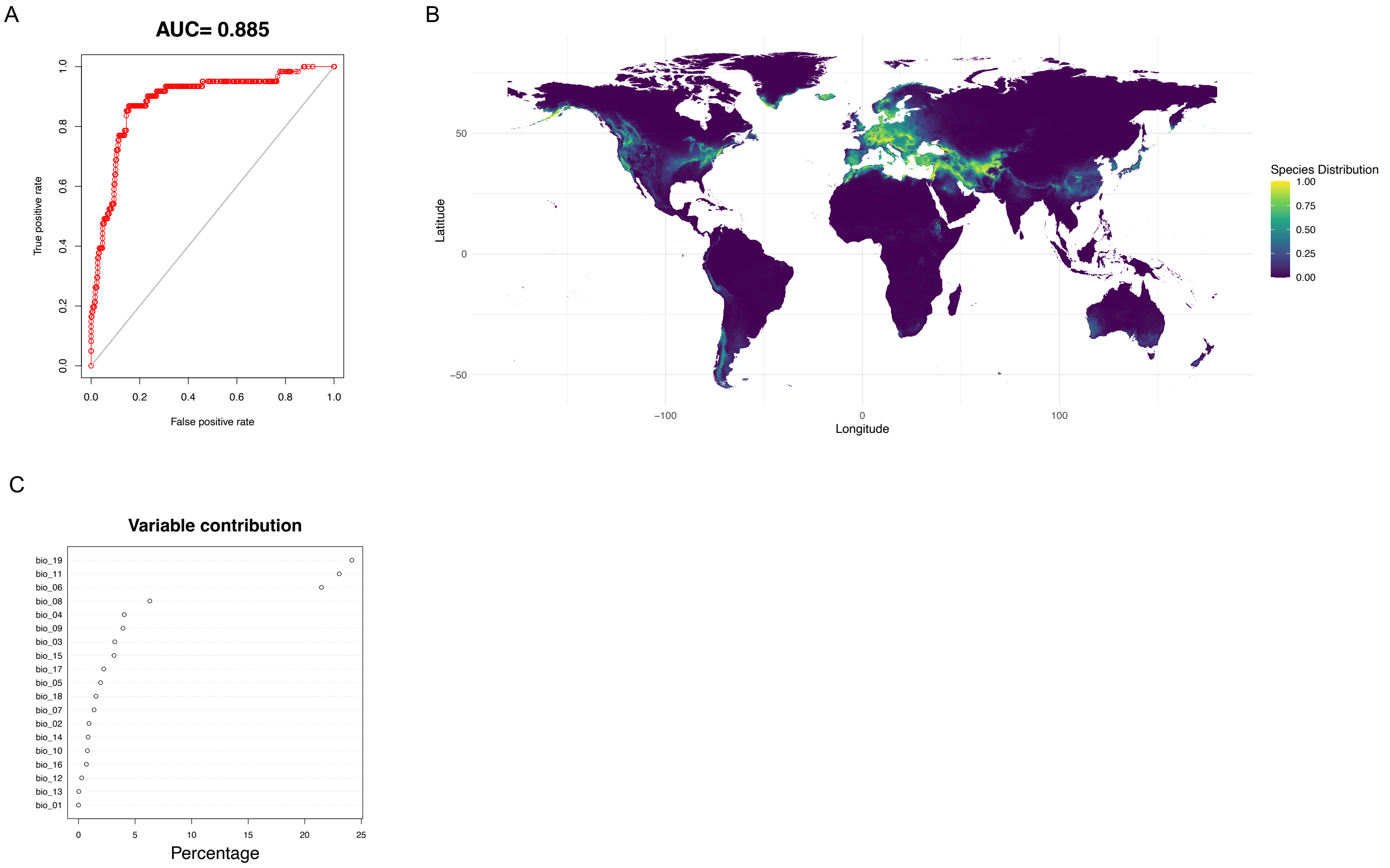


***Figure S7.*** *Species distribution modelling results for population 2* ***(A)*** *Area under the curve (AUC) graph* ***(B)*** *Species distribution map and* ***(C)*** *Variable contribution graph for the WorldClim bioclimatic variables examined.*

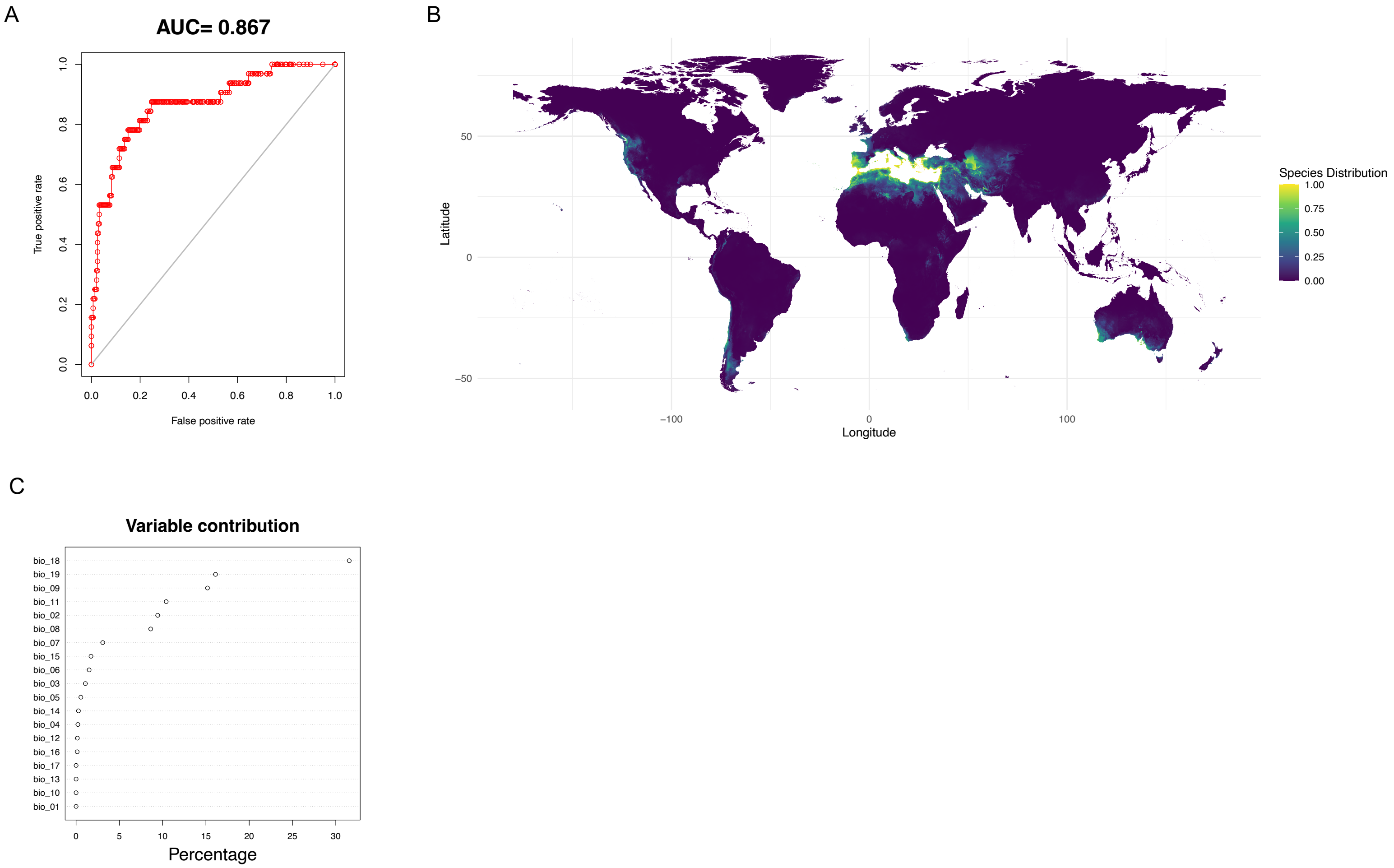


***Figure S8.*** *Species distribution modelling results for population 3* ***(A)*** *Area under the curve (AUC) graph* ***(B)*** *Species distribution map and* ***(C)*** *Variable contribution graph for the WorldClim bioclimatic variables examined.*

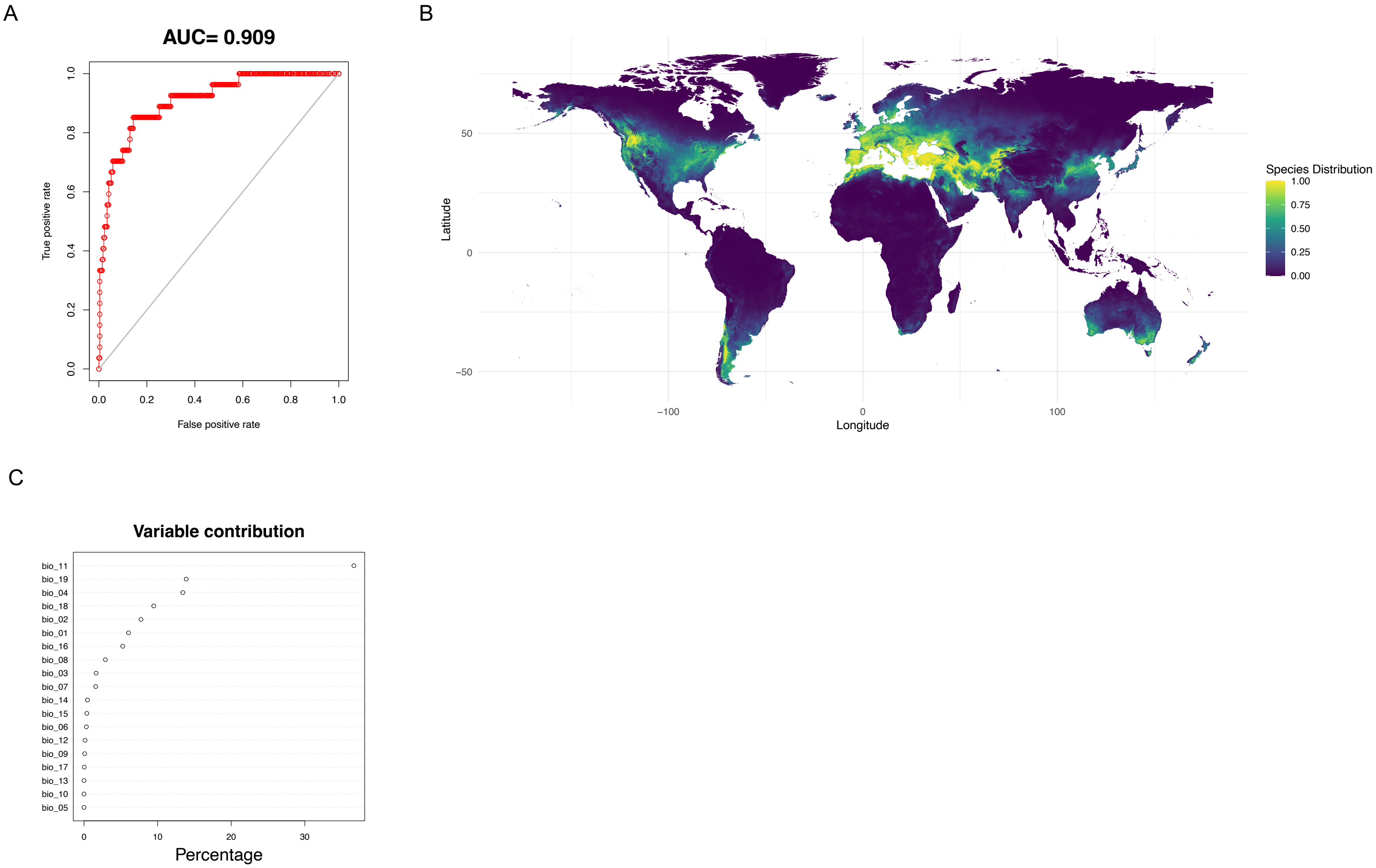


***Figure S9.*** *Species distribution modelling results for population 4* ***(A)*** *Area under the curve (AUC) graph* ***(B)*** *Species distribution map and* ***(C)*** *Variable contribution graph for the WorldClim bioclimatic variables examined.*

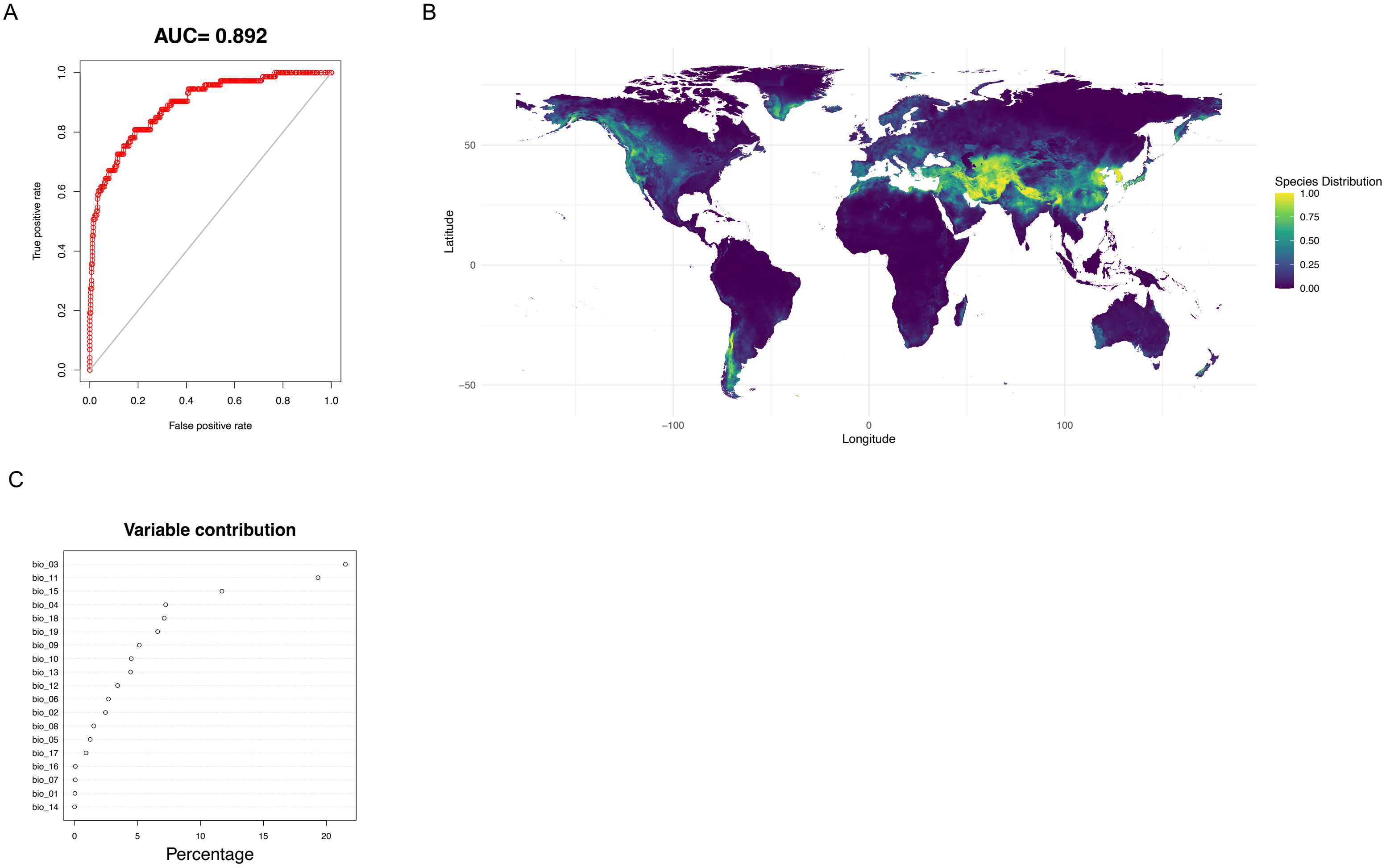


***Figure S10.*** *Species distribution modelling results for population 5* ***(A)*** *Area under the curve (AUC) graph* ***(B)*** *Species distribution map and* ***(C)*** *Variable contribution graph for the WorldClim bioclimatic variables examined.*

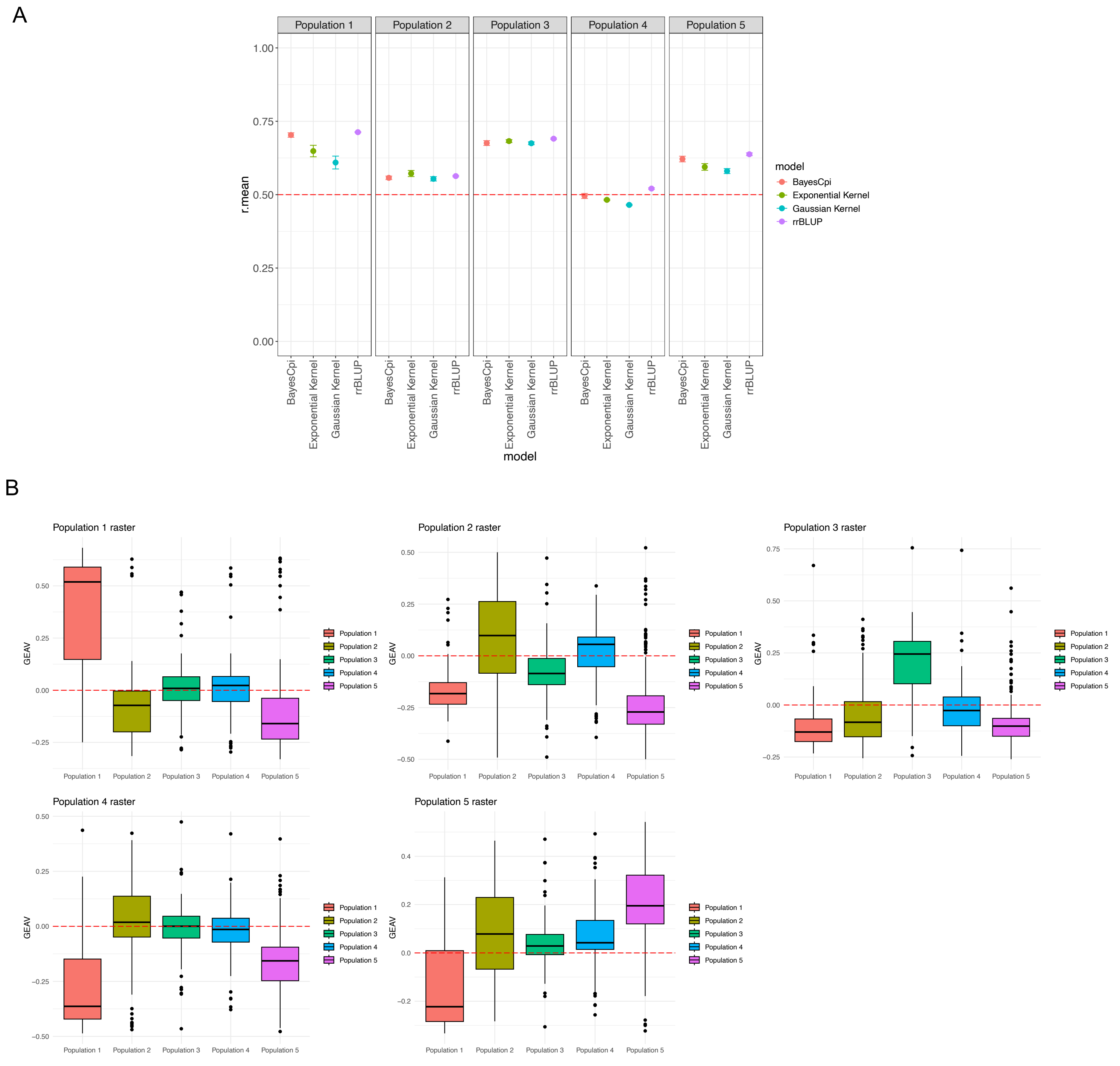


***Figure S11****. Prediction accuracy for Species Distribution Scores* ***(A)*** *using four genomic prediction methods, rrBLUP, G-BLUP with an exponential kernel, G-BLUP with a Gaussian Kernel and BayesCpi at k-fold = 10. Training set = 31, test set = 753* ***(B)*** *GEAVs by population using SDM suitability scores.*

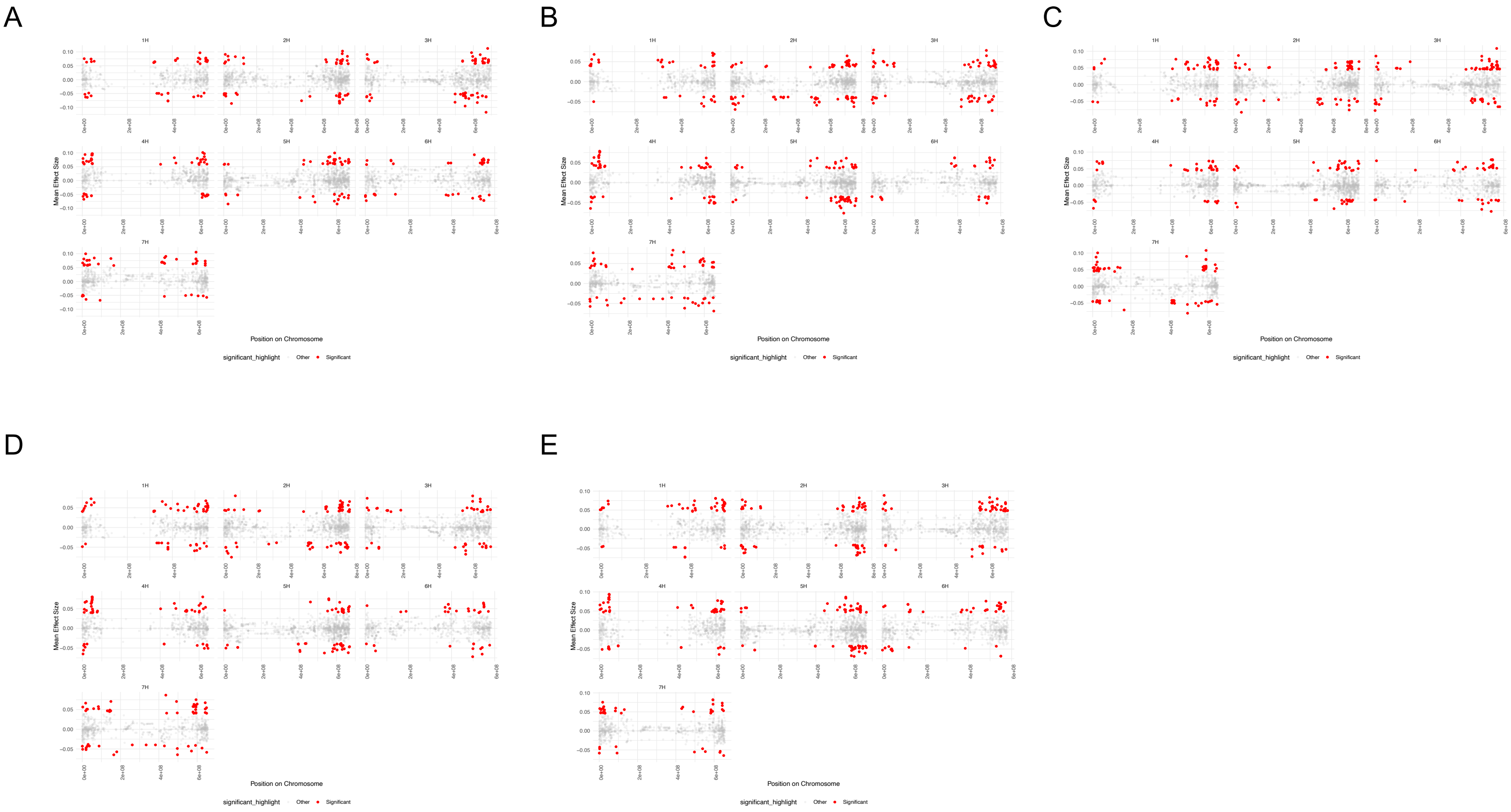


***Figure S12.*** *SNP effects by chromosome and populations. Mean SNP effects across all individuals in a population were calculated with SNP outliers identified by calculating z-scores (>3)* ***(A)*** *population 1 (363 outlier SNPs)* ***(B)*** *Population 2 (392 outlier SNPs)* ***(C)*** *population 3 (390 outlier SNPs)* ***(D)*** *Population 4 (384 outlier SNPs)* ***(E)*** *population 5 (372 outlier SNPs).*

A

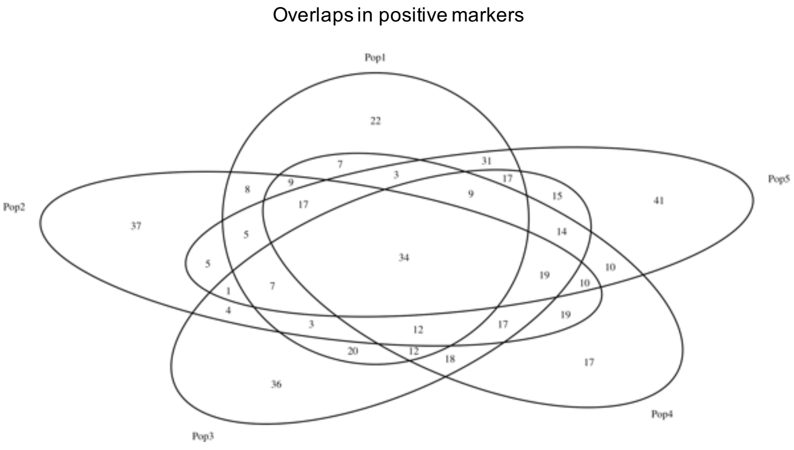


B

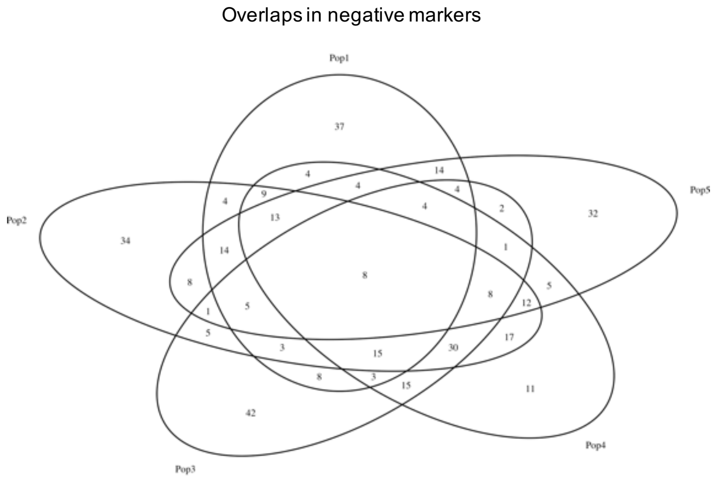


***Figure S13.*** *Overlaps in highest z-score SNPs across populations* ***(A)*** *positive effect SNP overlaps* ***(B)*** *negative effect SNP overlaps.*

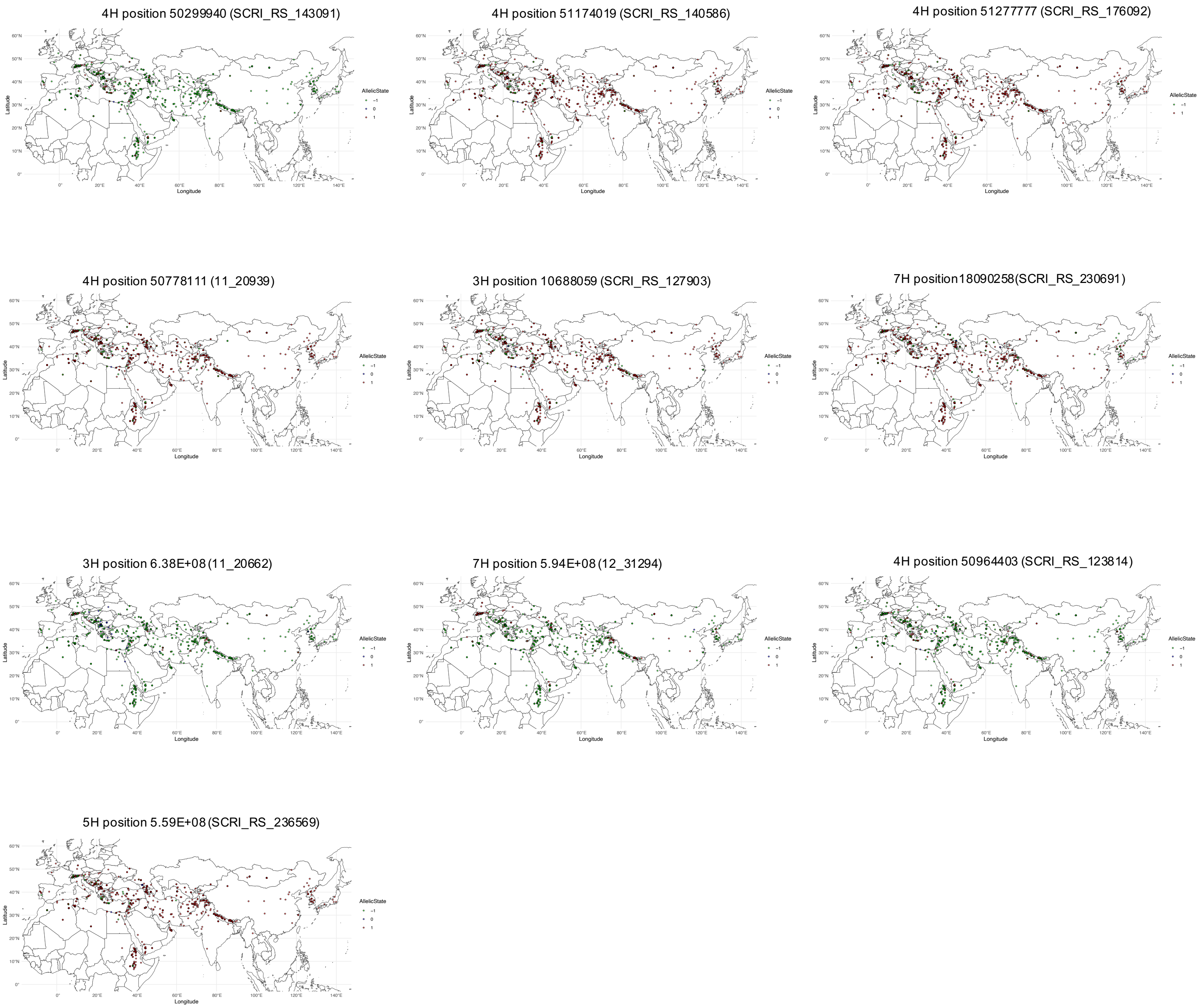


***Figure S14***. *Geographic distribution of allele variation for the top ten SNPs with the highest mean effect for Bio1.*

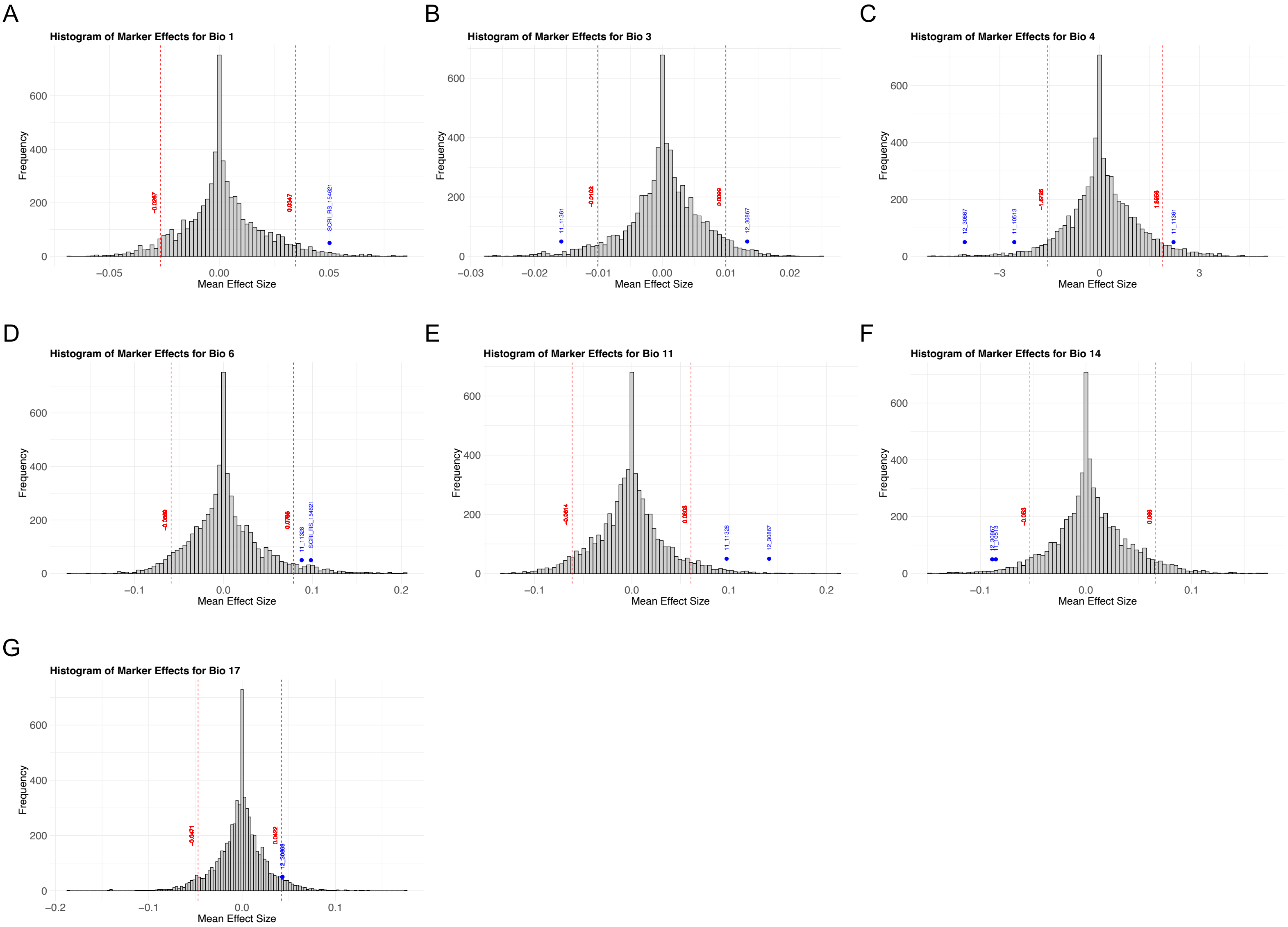


***Figure S15.*** *Histograms of marker effects for environmental variables with famous genes indicated in blue and the 5th and 95th percentiles indicated by a red dashed line **(A)** Bio 1 - Annual mean temperature **(B)** Bio 3 - Isothermality **(C)** Bio 4 - Temperature seasonality **(D)** Bio 6 - Min temperature of the coldest month **(E)** Bio 11 - Mean temperature of the coldest month **(F)** Bio 14 - Precipitation of the driest month **(G)** Bio 17 - Precipitation of the driest quarter.*

A

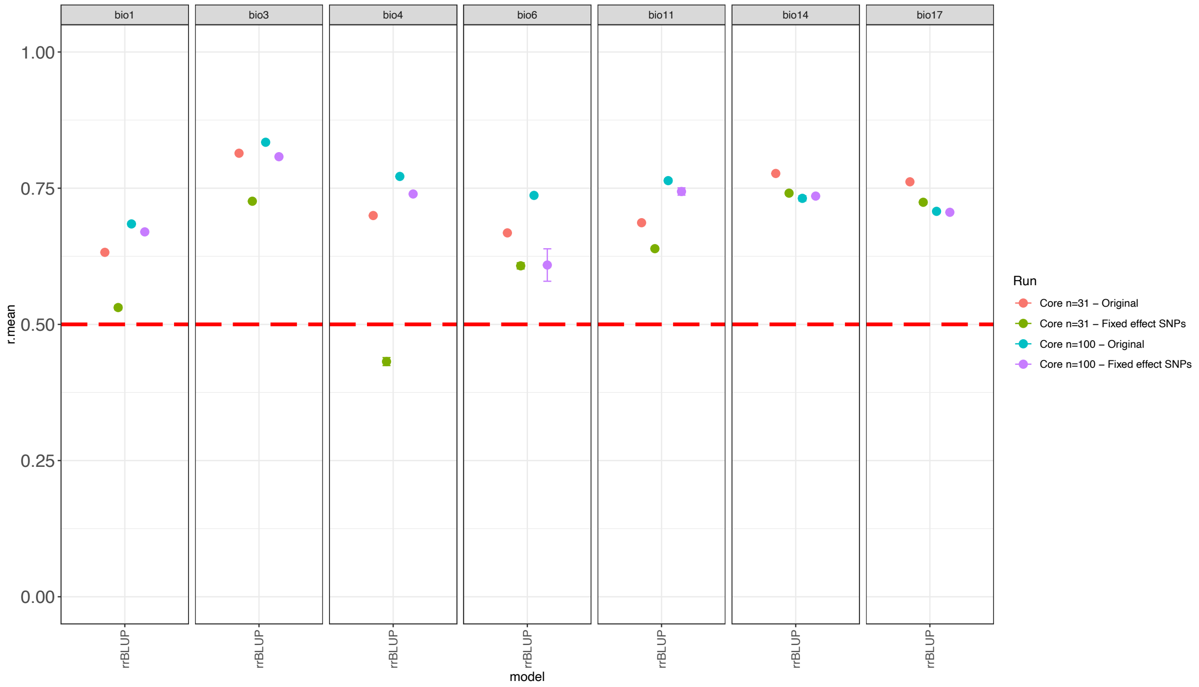


B

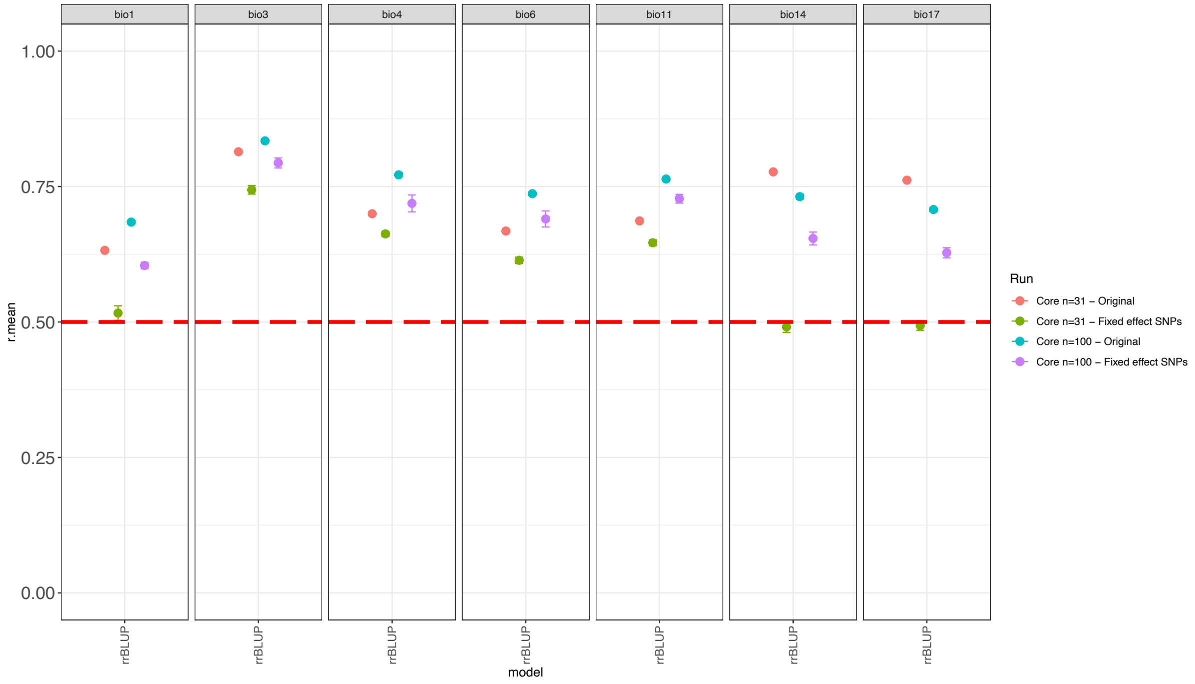


***Figure S16.*** *Comparison of rrBLUP model accuracy for core collections (n=31 and n=100) with major effect famous gene markers as fixed effects (Fixed effect SNPs) versus all random markers (Original) for* ***(A)*** *cold tolerance related SNPs (n=5)* ***(B)*** *Flowering time related SNPs (n=9).*

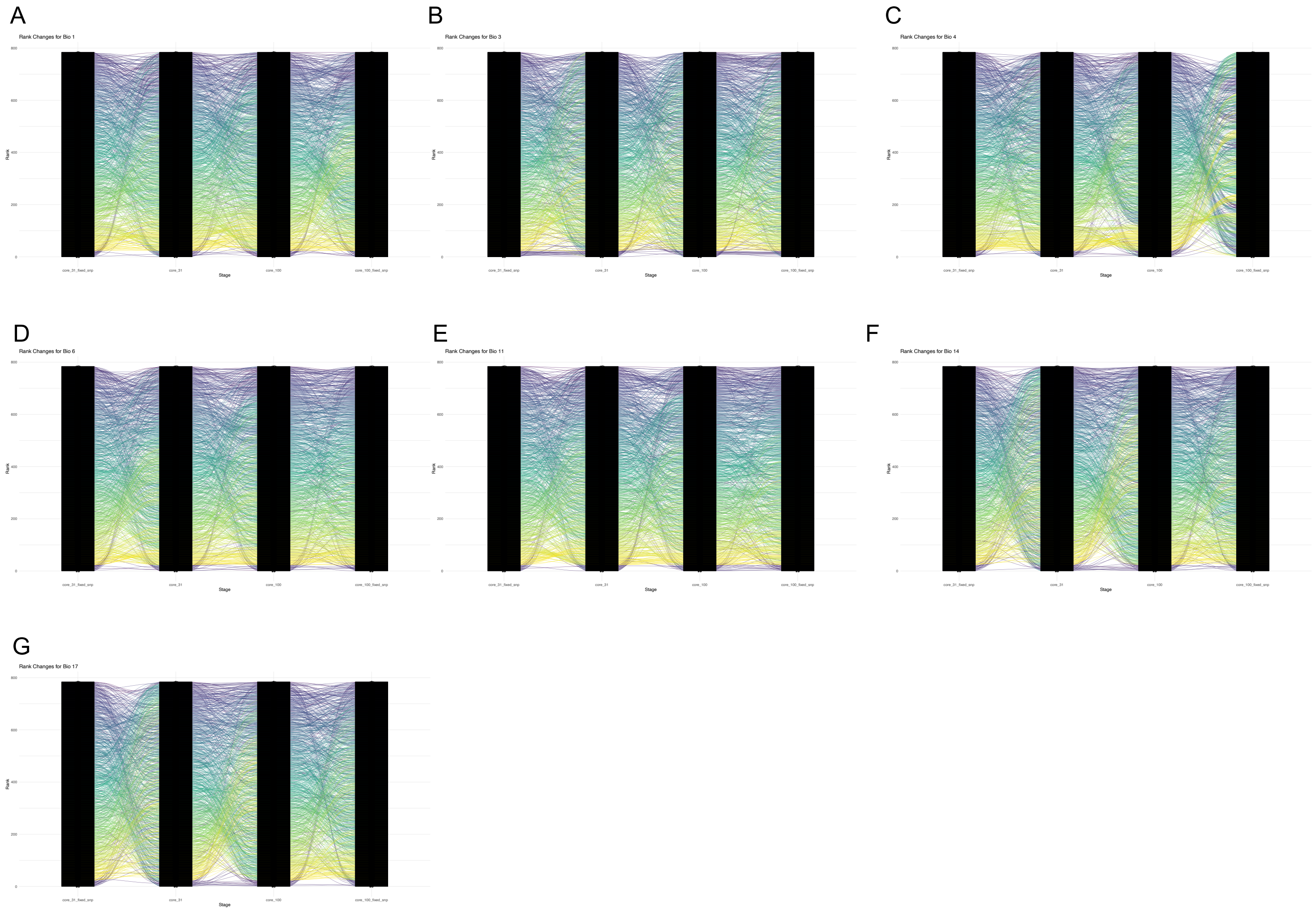


***Figure S17.*** *Comparing rank changes between cores (n=31 and n=100) with all markers as random effects and with fixed effect (famous genes n=14) SNPs* ***(A)*** *Bio 1 - Annual mean temperature* ***(B)*** *Bio 3 - Isothermality* ***(C)*** *Bio 4 - Temperature seasonality* ***(D)*** *Bio 6 - Min temperature of the coldest month* ***(E)*** *Bio 11 - Mean temperature of the coldest month* ***(F)*** *Bio 14 - Precipitation of the driest month* ***(G)*** *Bio 17 - Precipitation of the driest quarter.*
